# PD-1 controls differentiation, survival, and TCR affinity evolution of stem-like CD8+ T cells

**DOI:** 10.1101/2024.08.02.606241

**Authors:** Jyh Liang Hor, Edward C. Schrom, Abigail Wong-Rolle, Luke Vistain, Wanjing Shang, Qiang Dong, Chen Zhao, Chengcheng Jin, Ronald N. Germain

## Abstract

Stem-like progenitors are a critical subset of cytotoxic T cells that self-renew and give rise to expanded populations of effector cells critical for successful checkpoint blockade immunotherapy. Emerging evidence suggests that the tumor-draining lymph nodes can support the continuous generation of these stem-like cells that replenish the tumor sites and act as a critical source of expanded effector populations, underlining the importance of understanding what factors promote and maintain activated T cells in the stem-like state. Using advanced 3D multiplex immunofluorescence imaging, here we identified antigen-presentation niches in tumor-draining lymph nodes that support the expansion, maintenance, and affinity evolution of a unique population of TCF-1+PD-1+SLAMF6^hi^ stem-like CD8+ T cells. Our results show that contrary to the prevailing view that persistent TCR signaling drives terminal effector differentiation, prolonged antigen engagement well beyond the initial priming phase sustained the proliferation and self-renewal of these stem-like T cells *in vivo*. The inhibitory PD-1 pathway plays a central role in this process by mediating the fine-tuning of TCR and co-stimulatory signal input that enables selective expansion of high affinity TCR stem-like clones, enabling them to act as a renewable source of high affinity effector cells. PD-1 checkpoint blockade disrupts this fine tuning of input signaling, leading to terminal differentiation to the effector state or death of the most avid anti-tumor stem-like cells. Our results thus reveal an unexpected relationship between TCR ligand affinity recognition, a key negative feedback regulatory loop, and T cell stemness programming. Furthermore, these findings raise questions about whether anti-PD-1 checkpoint blockade during cancer immunotherapy provides a short-term anti-tumor effect that comes at the cost of diminishing efficacy due to progressive loss of these critical high affinity stem-like precursors.

## Main

Inhibitory molecules acting in *cis* (within a T cell) and *trans* (between cells) are central to the complex regulatory circuit that fine-tunes the functional output of T cell receptor (TCR) signaling. Programmed cell death 1 molecule (PD-1) and its ligands, PD-L1 and PD-L2, represent a key trans-acting inhibitory pathway that attenuates lymphocyte activation, restrains effector differentiation, and limits pathology in autoimmune diseases and chronic infections^1^. These checkpoint inhibitory pathways are now of substantial clinical interest due to their demonstrated capacity to promote anti-tumor T cell responses^2^. However, only a subset of patients respond favorably to checkpoint immunotherapy and many patients eventually experience acquired resistance to treatment and tumor relapse^3^.

Stem-like progenitor CD8+ T cells (T_SL_) represent a subset of activated cytotoxic T lymphocytes that retains high proliferative potential and self-renewal capacity^4^. Key studies have identified this subset or their immediate progeny as primarily responsible for the proliferative burst that gives rise to a large pool of functional effector cells during anti-PD-1 checkpoint immunotherapy in chronic viral infections and tumor-bearing hosts^5,6^. Emerging evidence suggests that tumor-draining lymph nodes (tdLNs) serve as reservoirs from which freshly generated T_SL_-like cells continuously replenish the tumor microenvironment (TME) and provide a source of functionally competent effector cells during checkpoint therapy^7-10^.

The questions of how activated T cells remain in a stem-like state in the lymphoid tissues during ongoing antigen stimulation and inflammation, especially under persistent antigen encounter in the case of tumors, and whether T cell fate is imprinted during early priming events or continues to be dynamically shaped by diverse inputs present in the LN microenvironment, remain unsettled. We employed 3D volumetric imaging techniques to approach these questions.

### PD-1+ SLAMF6+ T_SL_ are enriched in late antigen presentation niches within the tdLN

To map the spatial distribution and functional states of tumor antigen-specific CD8+ T cells in the tdLN, we performed 3D tissue imaging^11,12^ on optically cleared Day 8 tdLN slices (∼300µm thickness, comprising ∼25-30% of the whole tdLN volume). These slices were derived from tdLN of mice adoptively transferred with a physiological frequency of naïve OT-I precursors (500 cells)^13^ and intradermally implanted with a Kras/P53 lung adenocarcinoma line^14^ engineered to express ovalbumin antigen (KP-OVA). Fluorescent Xcr1 reporter mice were used as recipients to visualize XCR1+ cDC1^15^. We developed and optimized a 3D tissue staining protocol as well as a distributed computational pipeline that enables efficient single-cell level analysis of large 3D volume imaging datasets comprising >1 million cells of interest (**Extended Data Fig. 1a, Supplementary Video 1**). This provides a much more complete view of the cell populations in the tdLN than single cell studies that often only examine 1/10 this number of cells, while also adding high resolution spatial information.

Day 8 tdLN represented the peak expansion and differentiation stage of the primary adaptive immune response, during which the small number of naïve antigen-specific OT-I precursors substantially expanded and extensively infiltrated the LN T cell zone, interfollicular regions, and medullary sinuses (**Fig. 1a**). Using a combination of labeled antibodies to visualize stem-like markers (TCF-1, SLAMF6), activation state-associated proteins (BATF), a key inhibitory molecule (PD-1), and a feature of proliferating cells (Ki-67), we mapped the spatial distribution of expanded OT-I cells to their respective functional states in the tdLN (**Fig. 1b-e**). A small subset of OT-I cells formed distinct clusters in close association with XCR1+ cDC1 in the T cell zone. These clustering T cells expressed high levels of stem-like (TCF-1 and SLAMF6) and activation (PD-1, BATF) markers, indicative of a stem-like subset (**Fig. 1b**).

**Figure 1.**
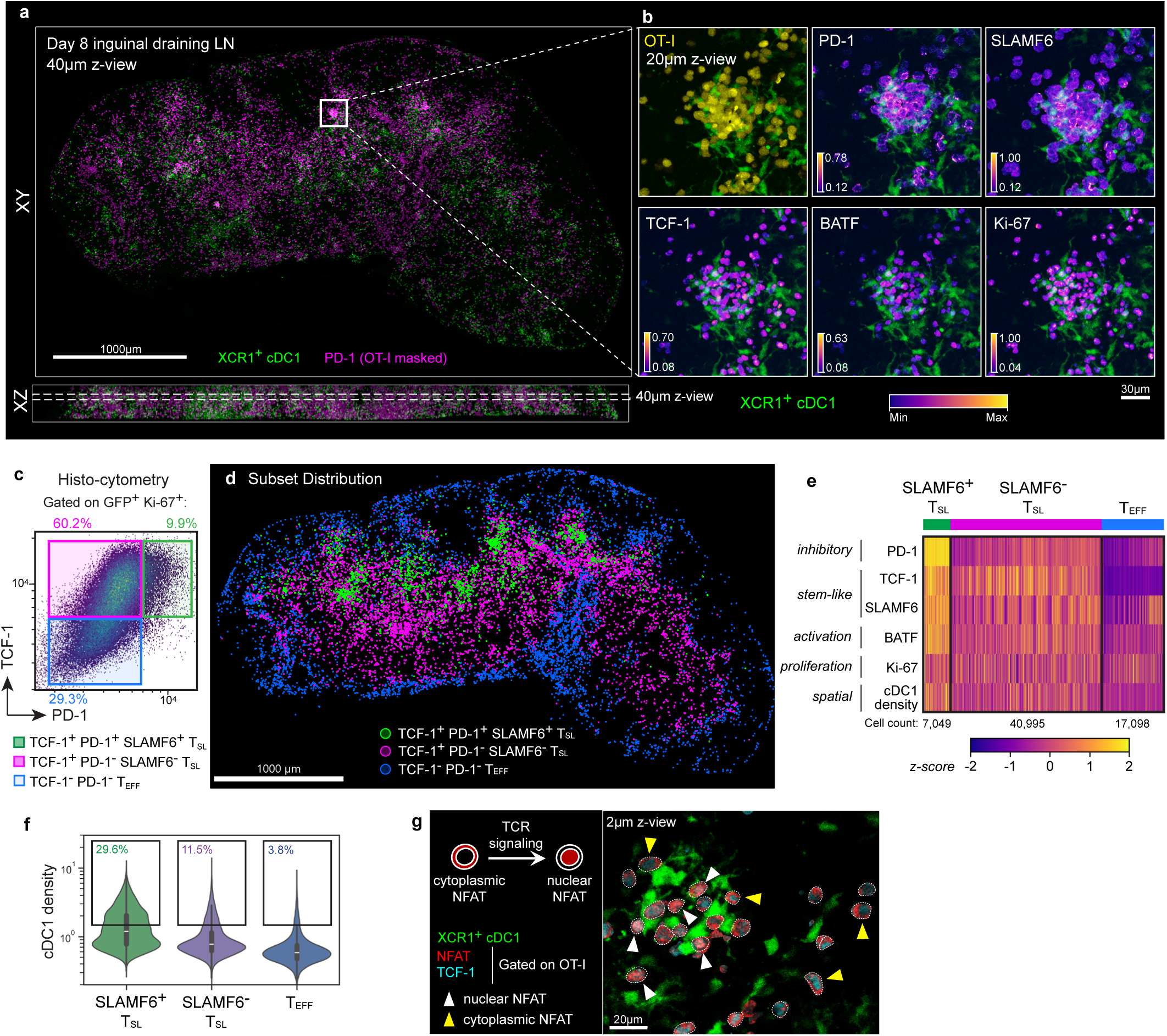
3D multiplexed tissue imaging revealed late antigen presentation niches in the tdLN. **a**, View of 40µm thick cross-section from a 300µm-thick optically cleared tissue slice from a Day 8 inguinal tdLN showing the spatial localization of expanded OT-I cells (color-coded in magenta based on PD-1 intensity) and XCR1+ cDC1 (green). White box indicates the close-up region displayed in (**b**). Mice were previously transferred with 500 naïve OT-I.GFP cells prior to tumor induction on the flank skin. **b**, Close-up images focusing on a cDC1-T clustering niche with the protein markers expressed by OT-I cells displayed with a perceptually uniform colormap, with min/max values scaled for each individual marker (1.0 = maximum bit-depth of image). **c**, Representative histo-cytometric gating strategy of three major OT-I subsets based on TCF-1 and PD-1 expression. **d**, Spatial distribution of OT-I subsets from (**a**) based on the gating strategy in (**c**). Cells within a 40µm z-thickness are displayed. **e**, Heatmap showing the normalized z-scores for each parameter of OT-I sorted by individual subsets. The heatmap contains the entire OT-I population of 65,142 cells. **f**, Quantification of OT-I subset proximity to dense cDC1 regions based on cDC1 spatial density (fluorescent XCR1 channel processed with a Gaussian filter with kernel of σ=3.27µm). T cells register higher cDC1 density value when positioned in denser cDC1 regions. Gates denote proportion of cells with cDC1 density values above the threshold of mean + 1 standard deviation, indicating close spatial proximity with cDC1. **g**, NFAT1 staining masked on OT-I cells from a close-up region in Day 8 tdLN, showing OT-I cells (dotted white outlines) clustering with cDC1 (green). White arrows indicate cells with nuclear NFAT localization (red) and their co-localization with nuclear TCF-1 stain (cyan). Yellow arrows show a few examples of OT-I cells with cytoplasmic NFAT localization. All data shown are representative of at least 3 independent experiments, with n=2-3 per experiment.

T-DC clustering is a classic feature of early T cell priming^16^. As naïve T cells also express high amount of TCF-1, we examined whether these clustering TCF-1+ cells belonged to a late infiltrating wave of naïve OT-I cells that had only recently experienced initial antigen exposure. Using either flow cytometry or imaging, we failed to detect residual proliferation dye signal among the antigen-specific OT-I cells in the tdLN after co-transfer of a mixture of pre-labelled naïve OT-I and polyclonal CD8+ T cells into tumor-bearing mice (**Extended Data Fig. 1b-f),** indicating that these clustering TCF-1+PD-1+SLAMF6+ cells at Day 8 represented a subset of activated and extensively divided OT-I T_SL_.

We then performed subset classification based on TCF-1 and PD-1 expression (**Extended Data Fig. 2a, Fig 1c-d, Supplementary Video 2**) with further quantitative analysis confirming that the TCF-1+PD-1+ subset is enriched with cells also showing high SLAMF6 and BATF expression and positioned in close proximity to dense cDC1 regions (**Fig. 1e, f, Extended Data Fig. 2b, c, Supplementary Video 3**). We refer to this TCF-1+ PD-1+ SLAMF6+ subset as ‘SLAMF6+ T_SL_’, and PD-1^int^ SLAMF6^int^ T_SL_ as ‘SLAMF6-T_SL_’. The protein expression signature of SLAMF6+ T_SL_ from these imaging data was also confirmed by flow cytometry (**Extended Data Fig. 2d, e**). The clear gradient of spatial distribution with respect to TCF-1 expression, with TCF-1+ T_SL_ preferentially localized to the T cell zone and TCF-1-effector cells (T_EFF_) in the LN periphery (**Fig. 1d, Supplementary Video 2**), conforms to previously reported observations^17^.

### SLAMF6+ T_SL_ are engaged in late antigen presentation with cDC1

As PD-1 upregulation is driven by TCR signaling, the pattern of elevated PD-1 (as well as BATF) expression among clustering SLAMF6+ T_SL_ suggested ongoing antigen signaling events occurring days beyond initial antigen contact. To examine if active TCR signaling was occurring within the niches containing SLAMF6+ T_SL_ and cDC1, we stained the tdLN tissues with antibodies against the transcription factor NFAT1, which translocates from the cytoplasm of T lymphocytes into the nucleus upon TCR engagement and NFAT dephosphorylation^18^ (**Fig. 1g**). We have previously shown that nuclear NFAT staining of T lymphocytes is highly specific for antigen-stimulated T cells in contact with antigen-presenting cells^19^. A high proportion of clustered OT-I T_SL_ spatially associated with cDC1 showed nuclear NFAT staining, whereas OT-I cells distant from cDC1 were largely devoid of this nuclear NFAT localization pattern (**Fig. 1g**). We also observed similar findings in the draining LNs from mice immunized with OVA protein and poly(I:C) as adjuvant (**Extended Data Fig. 2f, g**), suggesting that this late antigen presentation phase is not unique to the tumor context, but likely a fundamental characteristic of the primary adaptive immune response involving activated CD8+ T cells exhibiting a stem-like phenotype. Overall, these findings show that days after the initiation of TCR signaling and extensive cell division, the most stem-like population of CD8+ T cells preferentially remains associated with antigen-presenting cDC1 and responds to such antigen display with active TCR-dependent signaling. Such results contrast with the prevailing view that high intensity, prolonged antigen signaling necessarily drives T cells to an effector state.

### Preferential retention of SLAMF6+ T_SL_ in the tumor antigen-draining LN

We next examined the spatial distribution of polyclonal activated CD8+ T cell subsets by employing the same 3D microscopy technique as in **Fig. 1** with optically cleared tdLN tissues from Day 8 tumor-bearing mice that did not receive adoptive transfer of transgenic OT-I cells. This volumetric imaging technique enabled the detection of relative rare cellular niches that are often missed with thin tissue cryosections. Using a combination of CD8b, TCF-1, PD-1, SLAMF6 and Ki-67 markers to delineate activated Ki-67+ CD8+ cells, we observed the formation of CD8+ TCF-1+ T_SL_ clusters scattered throughout T cell zone, albeit at a much lower frequency and with a smaller cluster size than in the transgenic model (**Fig. 2a-c, Extended Data Fig 3a, b**). These clustering polyclonal CD8+ T cells also exhibited high level of PD-1 and SLAMF6 expression, similar to what was observed with OT-I. A density map of activated Ki-67+ CD8+ T cells revealed enrichment of staining for PD-1 and SLAMF6 in association with cDC1 within discrete foci of the tdLN (**Fig. 2a, Extended Data Fig. 3a**). Similar observations were also made examining the tdLN of MC38-OVA tumor-bearing mice, though with a lower frequency of T_SL_ clusters (**Extended Data Fig. 3c**). Overall, we noted that clustering SLAMF6+ T_SL_ niches is a distinct spatial feature that occurs even with natural precursor frequencies and with different tumor and vaccine models.

**Figure 2.**
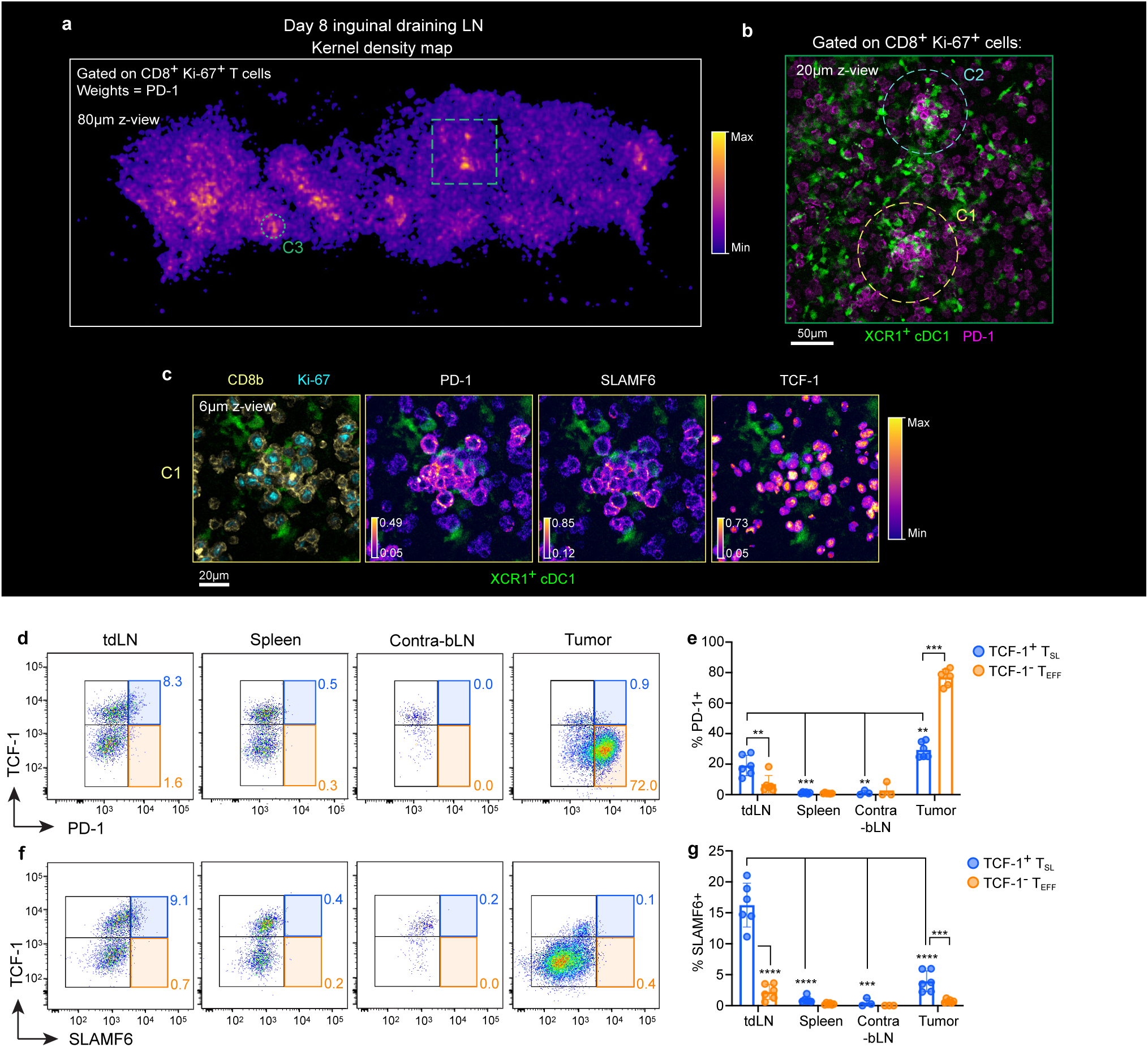
PD-1^+^ SLAMF6^+^ T_SL_ are uniquely retained in the tdLN. **(a-c)** Optically cleared tdLN slice from a Day 8 tumor-bearing mouse without adoptive transfer of transgenic T cells. Activated polyclonal CD8+ T cells are gated based on co-expression of CD8b and Ki-67. **a**, Kernel density map showing foci of activated polyclonal CD8+ T cells. Density was estimated using a Gaussian kernel of 6µm bandwidth, weighted on normalized PD-1 expression. Green box indicates close-up region as shown in (**b**), in which T-cDC1 clusters are further enlarged and displayed with individual protein markers: C1 in (**c**) and C2 in **Extended Data Fig. 2b**. Data shown are representative of two independent experiments (n=3 per experiment). (**d, f**) Representative gating strategy of OVA-tetramer+ CD8+ T cells recovered from tissues harvested from Day 10 tumor-bearing mice and analyzed using flow cytometry, based on TCF-1 and PD-1 (**d**) or SLAMF6 (**f**). The proportions of PD-1+ (**e**) and SLAMF6+ (**g**) of TCF-1+ T_SL_ and TCF-1-T_EFF_ subsets are quantified in (**e, g**). Data are pooled from 2 independent experiments (n=3-6). Error bars: means ± s.d. **p<0.01, ***p<0.001, ****p<0.0001; unpaired two-sided *t*-test.

These observations prompted us to ask if the SLAMF6+ T_SL_ subset was uniquely localized to the antigen-rich tdLN or also trafficked to other sites, including the TME. We employed flow cytometry and H-2K^b^-SIINFEKL tetramer staining to detect OVA-specific CD8+ T cells in the tumor-draining inguinal LNs, non-draining contra-lateral brachial LNs, the spleens and tumors at Day 10 post-tumor induction. While TCF-1+ T_SL_ and TCF-1-T_EFF_ were readily identified in all tissues examined, PD1^hi^ SLAMF6^hi^ T_SL_ were exclusively detected in the tdLN (**Fig. 2d-g**). As expected, PD-1 and SLAMF6 expression is highly correlated among TCF-1+ T_SL_ in the tdLN (**Extended Data Fig. 4a**). While SLAMF6 expression was elevated in all subsets of activated CD8+ T cells, PD1^hi^ T_SL_ had the highest SLAMF6 expression among all subsets, with T_SL_ in the tdLN showing a 3.5-fold greater expression compared to naïve CD8+ T cells (**Extended Data Fig. 4b, c**). Notably, unlike in the tdLN, PD-1+ T_SL_ in the tumor did not show high expression of SLAMF6 (**Fig. 2f, g, Extended Data Fig. 4b**). We found that SLAMF6+ T_SL_ subset also expressed higher amount of anti-apoptotic protein Bcl-2 than the other subsets (**Extended Data Fig. 4d**) and has enriched expression of the inhibitory molecule CD200 (**Extended Data Fig. 4e**), which has previously been identified as a marker associated with stem-like cells^7,20,21^.

High PD-1 expression outside of the tdLN was only observed in the tumor (**Fig. 2d, e**), in agreement with published findings^22^. T_EFF_ in non-tumor draining LN had low PD-1 levels, consistent with the notion that PD-1^hi^ T_SL_ involved in responses to late, ongoing antigen presentation were preferentially retained in the tdLN, while PD-1^lo^ T_SL_ and T_EFF_ that had egressed from the tdLN did not encounter cognate antigen-presenting cells until arriving at the tumor sites.

### Late antigen presentation is required for sustained expansion of CD8+ T_SL_

We next explored the functional role of late antigen presentation within tdLN. The XCR1-DTR transgenic mouse enables selective ablation of cross-presenting cDC1 using diphtheria toxin (DT) administration at various times after tumor implantation^15^. Our initial assessment of DT dosage determined that 2 successive doses are required to achieve >90% depletion in the LN (**Extended Data Fig. 4f-g**). Interestingly, even with very few cDC1 remaining upon DT treatment, we could still detect rare PD-1+ T_SL_ OT-I clusters formed around these cDC1 (**Extended Data Fig. 4h**).

Next, we initiated cDC1 depletion in KP-OVA tumor-bearing mice beginning at Day 5 post-induction (**Fig. 3a**), which represented the time point at which late antigen presentation clusters were first observed (data not shown). Using H-2K^b^-SIINFEKL tetramers to detect the OVA-specific CD8+ T cell population by flow cytometry at Day 10.5 (5.5 days post-depletion), we found that late cDC1 depletion resulted in significantly reduced OVA-specific CD8+ T cell expansion in the tdLN (**Fig. 3b**). Nearly all of the reduction was accounted for by a decline in the TCF-1+ T_SL_ compartment (**Fig. 3c**). Gating on the SLAMF6+ T_SL_ subset similarly revealed a marked reduction compared to non-depleted mice (**Extended Data Fig. 4i**, **Fig. 3d**). The T_SL_ compartment of cDC1-depleted tdLN also had a substantially diminished PD-1^hi^ subset (**Extended Data Fig. 4j**), with the mean PD-1 intensity of T_SL_ decreasing to that of their T_EFF_ counterparts in the tdLN (**Extended Data Fig. 4k**). This is consistent with the notion that ablation of antigen-presenting cDC1 preferentially eliminated late T_SL_ antigen signaling. In contrast, T_EFF_ expansion was only marginally affected by late cDC1 depletion in both the tdLN and the spleen (**Fig. 3c, Extended Data Fig. 4l**), with negligible effect in the tumor (**Extended Data Fig. 4m**). These experiments revealed that late antigen presentation by cDC1 in the tdLN is necessary for maintaining a large population of antigen-specific SLAMF6+ T_SL_.

**Figure 3.**
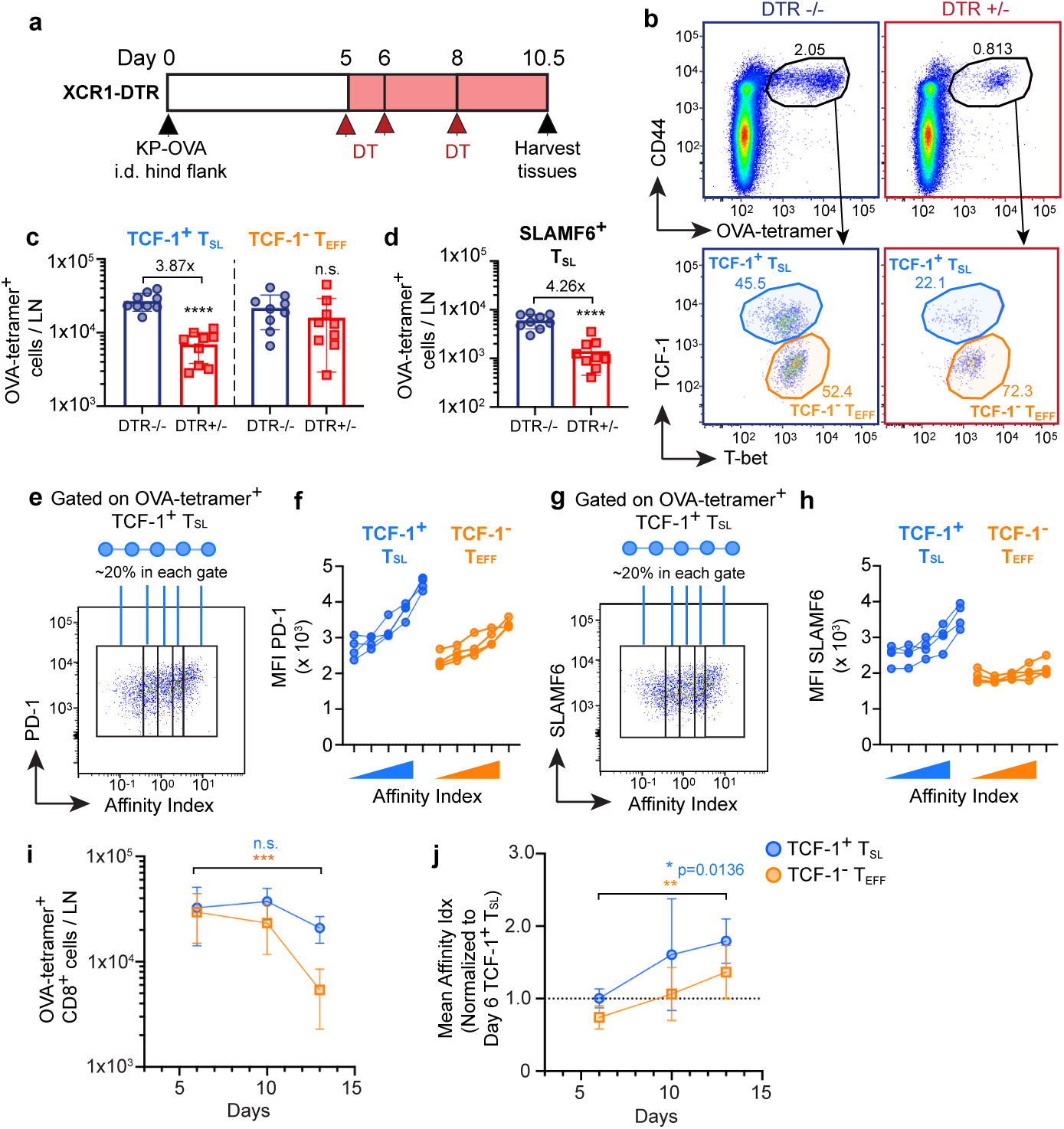
Antigen-driven expansion and TCR affinity evolution of TCF-1+ T_SL_ in the tdLN. **a**, Experimental scheme illustrating cDC1 depletion strategy in KP-OVA-bearing XCR1-DTR transgenic mice. **b**, Representative flow cytometry plot showing the gating strategy for OVA-tetramer+ CD8+ T cells (top) from Day 10 tdLN and further gated on TCF-1+ T_SL_ and TCF-1-T_EFF_ subsets (bottom). **c, d,** Quantification of OVA-specific TCF-1+ T_SL_ and TCF-1-T_EFF_ (**c**) as well as TCF-1+ SLAMF6+ T_SL_ (**d**) recovered from the tdLN. Data from 3 independent experiments (n=9 per group). (**e, g**) Representative flow cytometry plots showing PD-1 (**e**) and SLAMF6 (**g**) expression against TCR affinity index (tetramer:CD3 staining ratio) of OVA-tetramer+ TCF-1+ T_SL_ from Day 10 tdLN, and the gating strategy that divides each population into five separate bins each comprising ∼20% of the total subset population. (**f, h**) Mean fluorescence intensity of PD-1 (**f**) and SLAMF6 (**h**) of the binned subsets from (**e**) and (**g**). Data are representative of 2 independent experiments (n=4 per experiment). **i**, Total number of OVA-tetramer+ CD8+ TCF-1+ T_SL_ and TCF-1-T_EFF_ cells recovered from the tdLN across different time points. **j**, TCR affinity index plot of OVA-tetramer+ TCF-1+ T_SL_ and TCF-1-T_EFF_ in the tdLN showing the evolution of average TCR affinity indices from Day 6 to Day 13 post-tumor induction. Affinity indices are normalized to the mean affinity index of Day 6 TCF-1+ T_SL_ of the same experiment. Data are pooled from 2 independent experiments (n=7-8 per time point). Error bars: means ± s.d. *p<0.05, **p<0.01, ***p<0.001, ****p<0.0001; unpaired two-tailed *t*-test (**c, d**) and one-way ANOVA with Tukey’s multiple comparisons (**i, j**).

### Selective enrichment of high affinity TCR among SLAMF6+ T_SL_ in the tdLN

An important question is whether the T cells that adopt the PD-1+SLAMF6+ T_SL_ fate in the tdLN are a functionally selected subset of all the antigen-reactive T cells, or if they are simply a cross-section of all such cells who by virtue of their positioning within T_SL_-cDC1 cellular niches have taken on this phenotype. In particular, we wished to determine if the TCR of a T cell dictated its fate^23^. To this end, we performed further analysis of H-2K^b^- SIINFEKL tetramer-stained cells and found that tetramer binding was strongly correlated with PD-1 expression. Because tetramer binding intensity alone can vary based on the total amount of TCR expressed on the cell surface, we normalized tetramer staining intensity to CD3 intensity (tetramer:CD3 staining ratio) to derive a “TCR affinity index” of the T cells (see Methods for a detailed explanation of and rationale for this approach). Next, because individual adult mice each possesses a unique T cell repertoire spanning a range of TCR affinities, we further sub-divided the OVA-specific CD8+ T cells from each mouse’s tdLN samples into 5 separate bins of affinity indices, with each containing 20% of the total OVA-tetramer+ T_SL_ population in that tdLN (**Fig. 3e**). We found that the compartment containing the highest TCR affinity index was also associated with the highest mean expression of PD-1 within the T_SL_ subset (**Fig. 3f**).

Similarly, SLAMF6 protein expression was also highly correlated with both PD-1 expression and the TCR affinity index of T_SL_ (**Fig. 3g, h**). Because SLAMF6+ T_SL_ express higher levels of the co-receptor CD8 known to enhance tetramer binding, compared to the other subsets (data not shown,^24^), we performed further analysis by gating on an intermediate band of CD8-expressing T_SL_ (CD8-int). PD-1 expression remained positively correlated with TCR affinity index even under such constraints (**Extended Fig. 5a**). Additionally, we observed a similar PD-1 and TCR affinity index correlation in the draining LN of mice immunized with OVA protein (**Extended Data Fig. 5b, c**). Importantly, the positive correlation between PD-1 and TCR affinity is seen uniquely in the antigen-draining LN and is not observed in the spleen and other non-antigen presenting tissues (**Extended Data Fig. 5c**). These data collectively suggest that in a polyclonal CD8+ T cell setting, T cells with higher TCR affinity are selectively enriched among cells adopting the PD-1+SLAMF6+ T_SL_ fate.

### Late antigen presentation niches drive affinity evolution of CD8+ T_SL_

The presence of distinct cellular niches in a physiological context suggests the possibility of a selection mechanism that can be biologically relevant. The combined observations from imaging that revealed enriched clusters of PD-1+SLAMF6+ polyclonal CD8+ T_SL_ around cDC1 (**Fig. 2a-c**), together with flow cytometry data showing a positive correlation of PD-1 and SLAMF6 expression with TCR affinity, suggest that higher affinity OVA-specific T_SL_ clones are predisposed toward occupying the late T_SL_-cDC1 antigen presentation niches.

We therefore hypothesize that these late antigen presentation niches drive affinity evolution of antigen-specific CD8+ T cell responses over time through selective enrichment and expansion of higher TCR affinity clones. Importantly, due to the precursor-product relationship between the less differentiated T_SL_ and the more terminally differentiated T_EFF_ subsets, such selective expansion of CD8+ T cell clones at the T_SL_ progenitor stage ensures the generation of both high affinity progenitors and eventually, amplification of their progeny as high affinity effector cells.

This model predicts a progressive increase in the average TCR affinity of both T_SL_ and T_EFF_ subsets in the tdLN over time. To test this hypothesis, we examined the TCR affinity indices of OVA-specific CD8+ T_SL_ and T_EFF_ from tdLNs collected at different time points after tumor induction. Indeed, despite the number of T_SL_ remaining relatively constant throughout, the average TCR affinity indices of both T_SL_ and T_EFF_ steadily increased over time from Day 6 to Day 13 (**Fig. 3i, j**). Further gating with respect to SLAMF6 expression revealed that such TCR affinity changes were particularly pronounced among SLAMF6+ T_SL_ (**Extended Data Fig. 5d-f**). Temporally, the TCR affinity index of T_EFF_ lagged slightly behind those of T_SL_, which agrees with the notion that the enrichment of higher affinity T_SL_ subsequently gave rise to the higher affinity T_EFF_. We also noted that extreme outliers can occur on occasion, which is likely caused by the random TCR repertoire generation that can sometimes give rise to a precursor population that skewed towards an extreme end of the affinity range^13^ (**Extended Data Fig. 5f**).

Altogether, our findings suggest that late antigen presentation is critical in facilitating the affinity evolution of CD8+ T_SL_ clones in a spatio-temporal manner.

### PD-1 signaling sustains and regulates the stem-like state of CD8+ T_SL_

The preceding results raise an important question, namely how high TCR affinity CD8+ T_SL_ retain their stemness while undergoing continued antigen receptor signaling and proliferation. Prior studies have established that TCR signaling strength is closely associated with CD8+ T cell differentiation fate, with stronger, more prolonged TCR signaling driving effector differentiation genes (e.g., T-bet and Blimp-1) whereas intermediate and weaker signaling promoting a memory phenotype (e.g., Bcl-6 and Eomes) (rev. in ^25^). These past observations would predict that sustained late antigen signaling by high affinity CD8+ T_SL_ should prime these cells toward terminal effector differentiation, in contrast to our observations.

Based on the elevated level of inhibitory PD-1 receptor expression among SLAMF6+ T_SL_, and some limited evidence that this molecule can contribute to preventing terminal differentiation of CD8+ T cells^26-28^, we surmised that PD-1 inhibitory function in the CD8+ T_SL_ could play a role in attenuating TCR and co-stimulatory signals, thus enabling the preservation of the stem-like state. We conducted high-resolution 3D imaging of tdLN slices stained with non-blocking anti-mouse PD-1 antibody (clone RMP1-30)^29,30^ to allow *in situ* detection of PD-1 engaged with its ligands. We found that clustered OT-I T_SL_ interacting with cDC1 demonstrated polarization of PD-1 towards the T-DC synaptic interface (**Fig. 4a**). These features resembled the formation of PD-1 microclusters previously reported to mediate inhibitory signaling through the recruitment of the phosphatase SHP2^31^. We included polyclonal anti-mouse PD-L1 antibody in subsequent tissue staining experiments, which revealed the co-localization of clustered PD-1 on OT-I and PD-L1 expressed on cDC1 (**Fig. 4b, Extended Data Fig. 6a**), indicating active engagement of PD-1 and its ligands during late antigen presentation.

**Figure 4.**
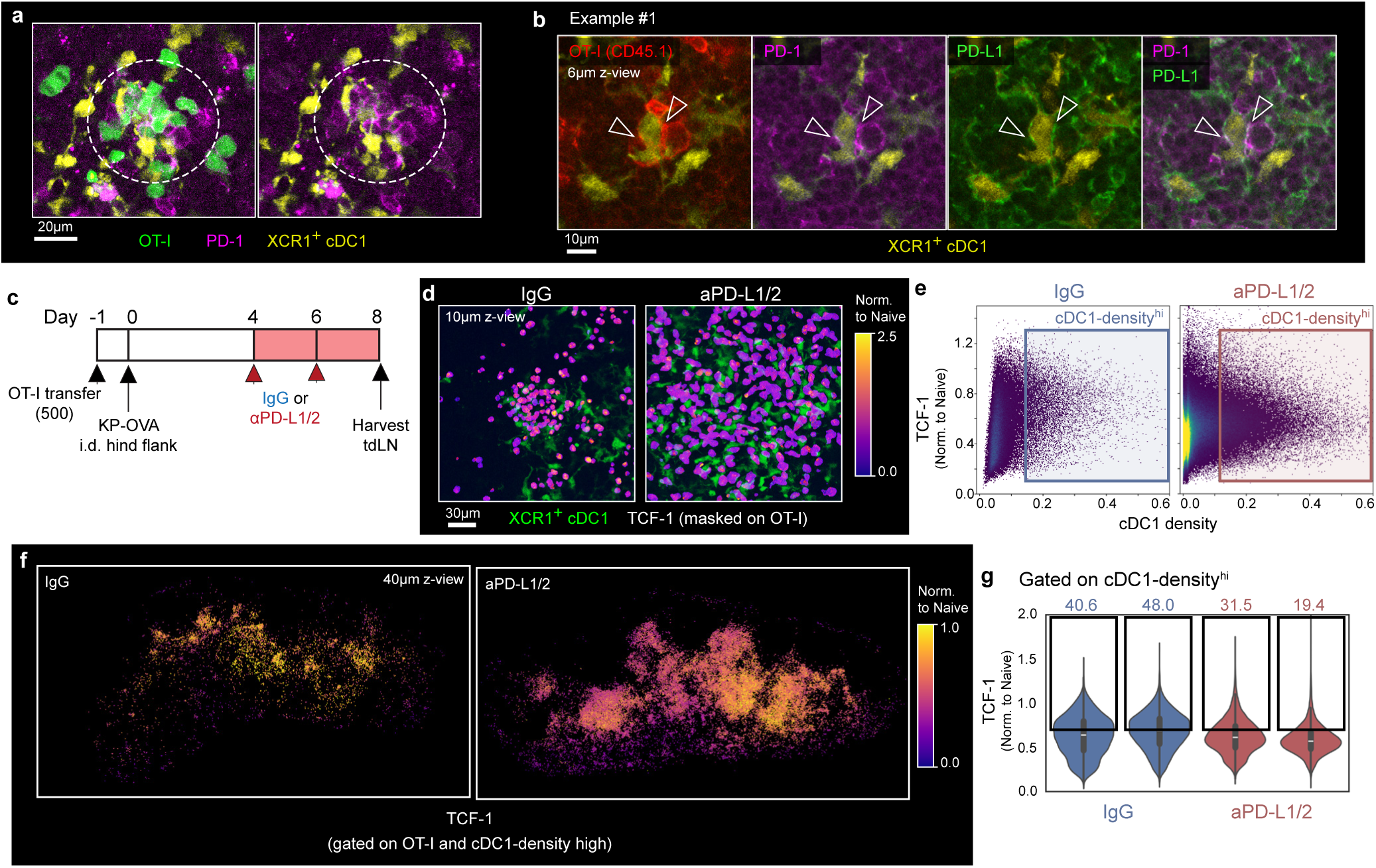
PD-1 signaling sustains and regulates stem-like state of CD8+ TCF-1+ T_SL_. **a**, High resolution 3D scans of cleared Day 8 tdLN stained with non-blocking anti-PD-1 antibody. A close-up example of TCF-1+ OT-I T_SL_ clustered around cDC1 network showing punctate microclusters of PD-1 proteins polarized toward T-DC synaptic interface. Right, image without OT-I.GFP channel. Data are representative of at least 2 independent experiments (n=2 per experiment). **b**, Representative images of an example of clustering OT-I cells and the co-localization of PD-1 (magenta) on OT-I.CD45.1 (red) with PD-L1 (green) expressed on cDC1 (yellow). Shown is a thin cross-section of a 3D stack (4µm z-thickness) and arrows indicate polarization and punctate microcluster formation of PD-1. Note the co-localization of PD-1 and PD-L1 blending into white pixels, indicating receptor-ligand engagement. Data are representative of at least 2 independent experiments (n=2-3 per experiment). **c**, Experimental scheme for (**d-g**) with tumor-bearing mice receiving either IgG (control) or anti-PD-L1/PD-L2 blocking antibodies from Day 4 onwards. **d**, Representative close-up images of TCF-1 expression (masked on OT-I cells) within a dense cDC1 region. **e**, Histo-cytometry plots showing the distribution of OT-I cells in Day 8 tdLN and their relative expression of TCF-1 (normalized to naïve T cells) and cDC1 density (proximity to dense cDC1 region). cDC1-dense^hi^ subset is thresholded on mean + 1 s.d. above log(cDC1 density) and their spatial distribution as well as relative TCF-1 expression are shown in (**f**) as a 40µm cross-section view of the tdLN. **g**, Quantification of TCF-1 expression among cDC1-dense^hi^ OT-I cells (e, f). Gates show the relative frequencies of cDC1-dense^hi^ OT-I with TCF-1 expression (normalized to naïve T cells) of >0.7. Data are representative of 2 independent experiments (n=2 per group per experiment).

Using flow cytometry to measure the co-stimulatory and co-inhibitory ligands expressed by DCs in the tdLN, we found that whereas PD-L1 was highly expressed among migratory DC subsets even at steady state, a dramatic increase in PD-L2 (another ligand of PD-1) can be observed in Day 8 tdLN, and especially among the cross-presenting cDC1 subset (**Extended Data Fig. 6b-g**). This suggests that both PD-1 ligands can potentially contribute to PD-1-mediated TCR signal attenuation during late antigen presentation.

To investigate the role of PD-1 inhibitory signaling in promoting the expansion/survival of T_SL_, we performed checkpoint co-blockade with anti-PD-L1 and anti-PD-L2 (**Fig. 4c**). tdLN harvested on Day 8 revealed a proliferative burst of OT-I after checkpoint blockade, as anticipated from the loss of PD-1 inhibition (**Fig. 4d**). Whereas OT-I spatially associated with the cDC1 network in the T cell zone displayed higher TCF-1 expression under control setting, an extensively divided OT-I population with heterogeneous expression of TCF-1 occupied the dense cDC1 region after checkpoint blockade (**Fig. 4d**). Quantitative analysis on the gated OT-I population in close spatial proximity to cDC1 showed that these T cells had lower mean TCF-1 expression than OT-I in control animals (**Fig. 4e-g**). Checkpoint blockade also yielded intensified cleaved caspase-3 staining across the tdLN, including OT-I cells that formed clusters with cDC1 (**Extended Data Fig. 7a, b**). Increased cleaved caspase-3 expression is associated with T cell contraction accompanied by apoptosis and cell death^32^. Strong antigen signaling especially in the absence of PD-1 inhibition is known to cause activated-induced cell death (AICD)^33^. Indeed, using flow cytometry to measure cleaved caspase-3 expression, OVA-tetramer+ SLAMF6+ T_SL_ in the tdLN showed significantly higher expression in anti-PD-L1/PD-L2 treated mice (**Extended Data Fig. 7c**). These data suggest that disruption of PD-1 signaling drives downregulation of TCF-1 among T cells receiving strong antigen signaling, together with an increase in apoptosis.

### PD-1 checkpoint blockade markedly reduces high affinity T_SL_ in the tdLN

Our findings so far suggest that blockade of PD-1 inhibitory signaling axis can disrupt maintenance of stem-like state and/or drive antigen-engaged T_SL_ towards cell death, potentially dysregulating the clonal landscape of antigen-specific CD8+ T cell repertoire. We therefore investigated the effects of PD-1 checkpoint blockade on polyclonal OVA-specific T cells from KP-OVA tumor-bearing mice, harvesting tdLN at Day 12 post-tumor induction (8 days after initial checkpoint blockade treatment) (**Fig. 5a**). Mice treated with a combined anti-PD-L1 and anti-PD-L2 from Day 4 showed robust proliferation of both OVA-specific T_SL_ and T_EFF_ as anticipated (**Fig. 5b, c, Extended Data Fig. 8c**), as well as partial suppression of tumor growth (**Extended Data Fig. 8a**). However, the proliferative bursts were accompanied by a phenotypic shift among the T_SL_ compartment, characterized by downregulation of stem-like markers (TCF-1 and SLAMF6) (**Fig. 5b, d, e**) and a decrease in SLAMF6+ T_SL_ numbers compared to non-treated tdLN (**Fig. 5c**).

**Figure 5.**
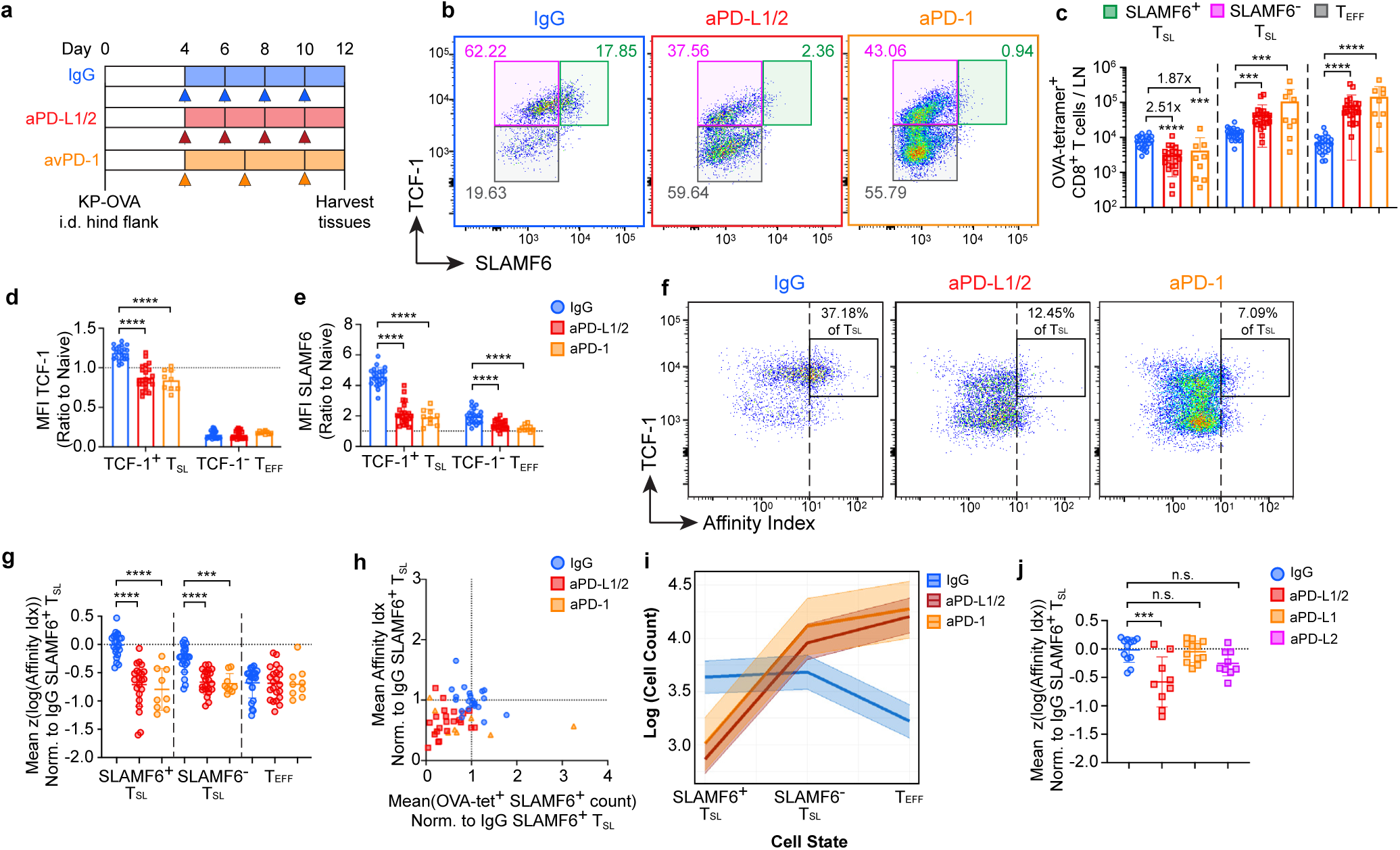
PD-1 checkpoint blockade disrupts accumulation and survival of high affinity CD8+ T_SL_. **a**, Experimental scheme illustrating checkpoint blockade treatment strategy. **b**, Representative flow cytometry plots of TCF-1 and SLAMF6 expression on OVA-tetramer+ CD8+ T cells from Day 12 tdLN from mice that had received either IgG (control), anti-PD-L1/PD-L2, or anti-PD-1 checkpoint blockade treatments. Gating strategy defines the three major T_SL_ and T_EFF_ subsets. The total numbers of each subset are given in (**c**). Mean fluorescence intensity of TCF-1 (**d**) and SLAMF6 (**e**) of each subset of OVA-tetramer+ CD8+ T cells, normalized to naïve (CD44-TCF-1+ PD-1-) CD8+ T cells. **f**, Representative flow cytometry plots showing TCF-1 expression against TCR affinity indices (tetramer:CD3 staining ratio) of OVA-tetramer+ CD8+ T cells of each treatment group. Box indicates a high affinity TCF-1+ T_SL_ sub-population based on an arbitrarily defined affinity index. Standardized log transformed TCR affinity indices of each treatment group are shown in (**g, j**). Each data point represents the mean value of subset from each animal. Affinity indices were normalized to the mean of log transformed affinity index of IgG (control) group SLAMF6+ T_SL_ from the same experimental cohort. **h**, Two-dimensional plot projecting the relationship between SLAMF6+ T_SL_ TCR affinity index versus SLAMF6+ T_SL_ cell count from each animal. TCR affinity index and cell count were normalized to the averages of IgG (control) group from the same experimental cohort. **i**, Bayesian multi-level linear modeling showing estimated high affinity (defined as “above average TCR affinity threshold”) cell count of each subset in logarithmic scale. Data from 7 independent experiments (n=3-4 per group per control group), with anti-PD-1 group from 2 independent experiments. Data from (**j**) are pooled from 3-4 independent experiments (n=3-5 per group per experiment). Error bars: means ± s.d. *p<0.05, **p<0.01, ***p<0.001, ****p<0.0001; one-way ANOVA with Tukey’s multiple comparisons.

As our findings also implicate PD-1 in controlling the expansion/persistence of high TCR affinity T_SL_ clones that would comprise the source of the most potent anti-tumor effectors, we also examined the effect of PD-1 checkpoint blockade on the T_SL_ clonal repertoire. Using our affinity index approach, we found that the average TCR affinity indices of the remaining SLAMF6+ T_SL_ and SLAMF6-T_SL_ were substantially lower after PD-1 checkpoint blockade (**Fig. 5f, g, Extended Data Fig. 8b**). Similar findings were also obtained when mice were treated with an anti-PD-1 blocking antibody clone that blocks interaction with both PD-L1 and PD-L2^34,35^ (**Fig. 5b-g, Extended Data Fig. 8a, b**), indicating that the observed phenotypic changes were directly related to PD-1 inhibitory signaling rather than the impaired function of its ligands PD-L1 and PD-L2, which can ligate with other proteins such as CD80^36,37^ and RGMb^38,39^ to mediate inhibitory functions. Due to a number of outliers observed (**Fig. 5h**), we conducted further analysis using Bayesian linear modeling, which agreed that the loss of high affinity SLAMF6+ T_SL_ was unlikely to be caused by outlier events (**Fig. 5i**, see **Methods** for details).

Because many factors influence tetramer binding capacity, we performed an additional control experiment in which mice transferred with monoclonal OT-I T cells (OT-I.Rag1) were subjected to the same anti-PD-L1/PD-L2 checkpoint treatment strategy. OT-I cells showed a similar decline in TCF-1 and SLAMF6 expression after checkpoint blockade (**Extended Data Fig. 8d-f**), but as expected for a monoclonal TCR population, the TCR affinity indices remained similar among cells from the different treatment groups (**Extended Data Fig. 8g, h**). Because PD-1 blockade also led to downregulation of CD8 co-receptor that can influence tetramer binding^24^, we also measured CD8 downregulation in SLAMF6+ T_SL_ OT-I from treated mice and confirmed that although OT-I experienced similar level of CD8 downregulation as cells in the polyclonal antigen-specific repertoire, their affinity indices remained unaffected (**Extended Data Fig. 8i**). Therefore, despite detecting some heterogeneity in tetramer staining, the tetramer binding differences revealed in mice with polyclonal repertoires cannot be attributed to these extrinsic changes caused by checkpoint blockade alone.

PD-1 is also highly expressed among activated CD4+ T cells, including both T helper cells and regulatory T cells (Treg). Both populations are known to have major influence on CD8+ T cell responses, including control of TCR affinity distribution^40^. We thus examined the contribution of CD4+ T cells to the observed changes in the differentiation states and clonal repertoire of CD8+ T cells after checkpoint blockade. Pre-depletion of both conventional CD4+ and Treg cells yielded enhanced SLAMF6+ T_SL_ generation in the tdLN (**Extended Data Fig. 9a, d-e**). Nonetheless, checkpoint blockade still resulted in significant downregulation of TCF-1 and SLAMF6 (**Extended Data Fig. 9a-c**), the dramatic loss of SLAMF6+ T_SL_ subset (**Extended Data Fig. 9d-e**), reduced TCR affinity indices (**Extended Data Fig. 9f-g**), as well as a similar partial inhibition of tumor growth as seen in non-CD4 depleted mice (**Extended Data Fig. 9h**). Therefore, CD4+ T cells did not appear to play a significant role in driving the loss of stem-like state and the change in clonal repertoire of CD8+ T_SL_ reported here.

Finally, given that our blocking strategy relied on the near complete repression of PD-1 mediated inhibition, we explored whether partial blockade in the form of single ligand antibody treatment (anti-PD-L1 or PD-L2 only) would be sufficient to reproduce the loss of high affinity CD8+ T_SL_ in the tdLN. We found that anti-PD-L1 blockade alone drove moderate but less effective tumor regression than co-blockade with PD-L1 and PD-L2 (**Extended Data Fig. 10a, b**), as well as a moderate expansion of both T_SL_ and T_EFF_ in the spleen (**Extended Data Fig. 10d-e**). Intriguingly, unlike with dual PD-1 ligand blockade, the numbers and TCR affinity index of SLAMF6+ T_SL_ after anti-PD-L1 blockade alone were not significantly lowered (**Fig. 5j, Extended Data Fig. 10c, f**). On the other hand, PD-L2 blockade alone did not induce significant expansion of either T_SL_ or T_EFF_, nor did it contribute to the control of tumor growth (**Extended Data Fig. 10a-d**).

Overall, these data reveal that strongly impaired PD-1 negative signaling is required to promote robust effector expansion capable of curbing tumor growth, but on the other hand, such thorough blockade of checkpoint molecules also resulted in the loss of high affinity SLAMF6+ T_SL_. These key precursors can be preserved with less effective interference with the PD-1 inhibitory axis, but which also substantially diminishes its effect on anti-tumor activity.

## Discussion

Contrary to the widely accepted model that stronger TCR signaling experienced by higher affinity cells drives terminal differentiation toward effector states^25^, we show here that prolonged antigen stimulation in the tdLN promotes the expansion of high TCR affinity clones as stem-like progenitors in a delicate process balanced by PD-1 inhibitory pathways. Antigen-bearing cDC1s serve as evolution sites that enable high affinity clones to receive TCR signals, proliferate and out-compete lower affinity clones over time. As T_SL_ are precursors to T_EFF_, expansion of high affinity T_SL_ subsequently gives rise to a large pool of high affinity T_EFF_. Such a mechanism is thus congruent with earlier studies showing strong effector bias among T cells with high TCR ligand affinity. However, it changes our understanding from one in which high affinity antigen recognition directly promotes exit from stem-like state, to one in which high affinity TCR signaling drives the enrichment of stem-like cells, whose progeny differentiate into effectors. Importantly, the latter enables high affinity clones to be preserved as T_SL_. The prolonged association of high affinity T_SL_ with cDC1 also helps explain prior observations that lower affinity antigen-specific clones egressed from lymphoid tissues earlier than their high affinity counterparts^41,42^. A recent study found that tumor antigen-specific T cells remained highly enriched in the tdLN as T_SL_ and differentiate into effectors upon migrating to the TME^43^. Another study reported that suboptimal TCR signaling in mouse tumor model promotes accelerated differentiation into effector cells^21^. Both reports are consistent with our findings here.

Our results implicate the PD-1 inhibitory pathway in maintaining stemness of high affinity T_SL_ through blunting of TCR input during prolonged antigen stimulation in the tdLN. While the locus of PD-1 inhibition^31,44,45^ has been debated, recent data from this laboratory confirm the TCR as the primary target^46^. This view is also supported by data showing a bias towards terminal differentiation of CD8+ T cells in PD-1 deficient mice^26,28^ and fits with a negative feedback model in which strong TCR signaling by high affinity T_SL_ drives strong upregulation of PD-1^27^. An earlier study proposed a similar affinity-biased model for control of effector CD4+ T cells, in which higher affinity T cells induce greater TCR-dependent CTLA-4 recruitment to the immune synapse, resulting in stronger inhibition of co-stimulatory signaling^47^. *In vivo* blockade with anti-PD-L1 prolongs effector CD4+ T-DC interactions, further supporting a key role for TCR-dependent PD-1 expression in regulating the duration and extent of high antigen level-dependent signaling^48^. These linked signaling and feedback mechanisms yield a finely tuned system controlling TCR input to support T_SL_ fate among an optimal cohort of T cells.

These findings raise important clinical implications regarding PD-1 checkpoint immunotherapy. Changes in the TIL clonal landscape, sometimes referred to as ‘clonal replacement’, have been reported in patients undergoing anti-PD-1 immunotherapy, with extinction of pre-existing clones and emergence of novel tumor-reactive clones^49-53^. Our pre-clinical model suggests that during PD-1 checkpoint blockade, inter-clonal competition from lower affinity clones as well as increased AICD among higher affinity clones could promote a repertoire shift. Such changes may compromise subsequent treatment efficacy, as higher TCR affinity/avidity CD8+ TILs are associated with better tumor clearance in anti-PD-1-treated patients^54-56^, as well as superior tumor infiltration and improved tumor control in adoptive cell therapy^57,58^. As high affinity CD8+ TILs can rapidly become dysfunctional upon exposure to persistent antigen in the TME, these cells may only have a narrow time window to facilitate effective tumor killing before losing functional cytotoxicity^42,59^. The capacity of non-metastatic tdLNs to act as an ongoing source of TCF-1+ T_SL_, as recently reported^7-9^, could become dysregulated by PD-1 checkpoint immunotherapy, leading to impaired long-term maintenance of high affinity T_SL_ clones in the tdLNs, the loss of which could be permanent due to reduced naïve T cell output from the adult thymus^60,61^. Recent clinical evidence reporting a declining efficacy of repetitive checkpoint treatment in patients experiencing tumor relapse fits with this view^62^. Finally, our findings indicate that partial blockade of PD-1 pathway could avoid loss of high affinity T_SL_ at the price of decreased anti-tumor efficacy in any single round of treatment, suggesting that a ‘sweet spot’ could be found through careful titration of checkpoint blockade antibody dosage.

In sum, our study provides insight into how PD-1 inhibitory pathway regulates finely tuned processes that enable the immune system to maximize the use of its diverse TCR repertoire without rapid loss of the most antigen-responsive cells. Interference with this tuning disturbs a carefully balanced system, with potentially detrimental long-term consequences to host responses. We propose that more careful attention to the counter-vailing effects of checkpoint blockade will be important in maximizing the clinical benefit of such treatments going forward.

## Acknowledgements

We thank T. Kaisho (RIKEN-Yokohama, and Osaka University, Japan) for the generous provision of transgenic mouse strains. We also thank P. L. Schwartzberg (NIAID, NIH) for helpful discussions. We would also like to thank all members of the Lymphocyte Biology Section for their feedback and support. We acknowledge the use of computational resources of the NIH HPC Biowulf cluster for imaging data processing and analysis.

## Author Contributions

J.L.H. and R.N.G. conceived the study and wrote the manuscript; J.L.H. designed and performed all experimental work, analyzed data, and developed computational pipelines for imaging analyses; E.C.S. designed and performed the TCR affinity modeling analyses; W.S., A.R-W., Q.D., C.Z., and C.J. generated tumor lines; L.V. performed custom dye conjugation of antibodies.

## Data and material availability

Python code generated for imaging data analysis are available at https://github.com/jlhor/3d-imaging-pipeline.

## Materials and Methods

### Mice

CD45.2 (C57BL/6J) and B6.GFP (C57BL/6-Tg(UBC-GFP)30Scha/J) mice were purchased from the Jackson Laboratory (strain numbers 000664 and 004354 respectively); CD45.1 (B6.SJL *Ptprc*^a^), OT-I.CD45.1 (B6(Ly5.1)-[Tg]TCR OT-I-[KO]RAG1), OT-I.CD45.2 (C57BL/6NAi-[Tg]TCR OT-1-[KO]RAG1) were obtained from the NIAID-Taconic exchange program (strain numbers 8478, 300 and 175, respectively). XCR1-DTR (B6.Cg-Xcr1^tm2(HBEGF/Venus)Ksho^) and XCR1-venus (B6.Cg-Xcr1^tm1Ksho^) transgenic mice^15^ were kind gifts from Tsuneyasu Kaisho (RIKEN-Yokohama and Osaka University). OT-I.GFP mice were cross-bred from OT-I.CD45.2 and B6.GFP and maintained as a homozygous strain in our laboratory. The majority of mice employed were female and aged 6-16 weeks at the beginning of experiments, with a small number of experiments performed in male mice. All mice were bred and maintained under specific pathogen-free conditions at an American Association for the Accreditation of Laboratory Animal Care (AAALAC)-accredited animal facility within NIAID and were used under study protocol LISB-4E approved by NIAID Animal Care and Use Committee (NIH).

### Tumor cell line generation

The KP-OVA tumor line was generated by lentiviral transduction of KP6-1B11 cells (a single colony subcloned from the KP1233 cell line derived from a Kras^G12D/+^ Trp53^-/-^ mouse^14^) with a lentivirus expressing full-length OVA (LRG-EFS-ZsGreen-P2A-OVA). pcDNA3-OVA (Addgene #64599) was cloned into the lentiviral vector LRG-EFS-ZsGreen-P2A, a plasmid modified from LRG vector^63^, a kind gift from Junwei Shi (University of Pennsylvania). ZsGreen+ KP6-1B11 cells were sorted into 96-well plates containing single cells per well. Single cell-derived clones with homogenous morphology and ZsGreen expression were selected and expanded for Western blot and flow cytometry validation using anti-OVA antibody (Invitrogen, PA1-196) and anti-mouse SIINFEKL-H-2K^b^ (clone 25-D1.16). The KP-OVA line was cultured and passaged in RPMI medium containing 10% fetal calf serum (FCS), L-glutamine (2 mM), penicillin (100 units/ml), streptomycin (0.1 mg/ml), sodium pyruvate (1mM), Hepes (10mM) and 2-mercaptoethanol (1mM) at 37°C/6.5% CO_2_.

The MC38-OVA tumor line was generated by lentiviral transfection of MC38 cells with lentivirus expressing full-length OVA (pLV-EF1a-OVA-puro). The MC38 cell line was generously provided by Martin Meier-Schellersheim (NIAID, NIH). pcDNA3-OVA (Addgene #64599) was cloned into the pLV-EF1a-puro lentiviral vector. Infected cells were selected with puromycin (8µg/ml) and OVA expression was assessed by flow cytometry using anti-mouse SIINFEKL-H-2K^b^ (Biolegend, clone 25-D1.16). The MC38-OVA line was cultured and passaged in DMEM medium containing 10% FCS, L-glutamine (2 mM), penicillin (100 units/ml), streptomycin (0.1 mg/ml), sodium pyruvate (1mM) and Hepes (10mM) at 37°C/6.5% CO_2_.

### Tumor induction and protein immunization

For tumor induction, mice were anesthetized with isoflurane inhalation (2% induction, 1-1.5% maintenance). Mice were shaved on the left flank with a Wahl clipper (Kent Scientific), depilated using Nair hair removal cream and the shaved flank washed thoroughly with water-soaked gauze. Tumor cells (KP-OVA and MC38-OVA) were harvested by washing with PBS, incubated with 0.25% Trypsin/EDTA (ThermoFisher Scientific) at 37°C for 5 min, then washed with pre-warmed RPMI supplemented with 10% FCS. 4×10^5^ KP-OVA or 5×10^6^ MC38-OVA were suspended in 20µl Hanks’ balanced saline solution (HBSS) and intradermally injected into the left hind flank skin using a 30G needle. Tumor volumes were measured using a digital caliper and the volume estimated using the formula: volume = ((width^2^ x length)/2). For OVA protein immunization, 25µg ovalbumin (OVA EndoFit, Invivogen) and 12.5µg poly(I:C) HMW (Invivogen) were reconstituted in 20µl PBS and were injected intradermally as above, or subcutaneously in the left footpad.

### Isolation of CD8+ T cells and adoptive cell transfer

Naïve OT-I or wildtype T cells (B6.GFP, CD45.1 or CD45.2) were isolated from spleens and lymph nodes of donor mice using magnetic cell separation (MACS) using mouse CD8a+ T cell isolation kits (Miltenyi Biotec), as per the manufacturer’s instructions. Between 500 and 2×10^3^ cells were injected intravenously in 200µl HBSS into recipient mice at least 1 day prior to tumor induction or immunization. For CellTrace Violet labeling, CellTrace Violet dye (ThermoFisher Scientific) was added at a final concentration of 5µM to 1×10^7^ cells/mL suspended in 0.1% bovine serum albumin-containing PBS and incubated at 37°C for 10 min, prior to addition of RPMI with 10% FCS to 10x staining volume and incubated further at 37°C for 5 min. Cells were washed in RPMI prior to resuspension in HBSS for adoptive transfer.

### In vivo antibody treatment

For blockade of PD-L1 and PD-L2, mice were intraperitoneally injected with 250µg anti-mouse PD-L1 (clone 10F.9G2; BioXCell) and anti-mouse PD-L2 (clone TY25; BioXCell) every 2 days starting on day 4 after tumor induction. For blockade of PD-1, 250µg anti-mouse PD-1 (clone 29F.1A12; BioXCell) was injected intraperitoneally into tumor-bearing mice every 3 days starting on day 4 post-tumor induction. Rat IgG2a (clone 2A3; BioXCell) and rat IgG2b (clone LTF-2; BioXCell) isotype antibodies were injected at a dose of 250µg each as control treatment. In some experiments, PBS was injected intraperitoneally into control animals. No phenotypic differences were observed between IgG-treated and PBS-treated controls.

For depletion of CD4 T cells, mice were intraperitoneally injected with 100µg anti-mouse CD4 monoclonal antibody (clone GK1.5; BioXCell) 3 and 1 day prior to tumor induction.

### XCR-DTR depletion

For diphtheria toxin (DT) depletion of cDC1 in XCR1-DTR mice, 1µg DT (Millipore Sigma, Cat. No: 322326) diluted in PBS was injected intraperitoneally into tumor-bearing mice on days 5, 6 and 8 after tumor induction.

### Tissue preparation for 3D imaging

Mice were euthanized with sodium pentobarbital through intraperitoneal injection, immediately followed by cardiac perfusion with 10ml of 1% paraformaldehyde (PFA) prepared from 16% aqueous stock (Electron Microscopy Sciences). Tissues were then harvested and fixed for 24 hours in 1ml fixative buffer (BD Cytofix/Cytoperm prepared 1:4 buffered to PBS; BD Biosciences) at 4°C on a slow shaker. Fixed tissues were then washed in 2ml PBS solution overnight and embedded in 4% UltraPure^TM^ low-melting point agarose (ThermoFisher Scientific) prepared with PBS (cooled to and maintained at 40°C in a water bath after boiling). Embedded tissues were allowed to solidify on ice. 300µm agarose-embedded tissue slices were cut using a VT1200S vibrating blade microtome (Leica) at a speed of 0.06mm/s and amplitude of 1.50. Tissue slices were collected into PBS-filled wells in 24-well plates and stored at 4°C.

### Antibody Conjugation

Custom antibody fluorophore labeling was performed when specific antibody-fluorophore pairs were not commercially available. Purified antibodies were concentrated using Amicon Ultra 50k MWCO centrifugal filters (Millipore). 10mM NHS ester-dye solution was prepared by dissolving NHS ester-dye with DMSO (Sigma-Aldrich). The NHS ester-dye solution and concentrated antibody solution were combined in a 1:9 ratio to yield a final NHS ester-dye concentration of 1mM, and solution was allowed to react on ice for 60 min. The reaction mixture was then diluted and concentrated in the centrifugal filters at least 3 times with PBS at 14,000 g centrifugation for 5 min to remove unbound dye. The final antibody-fluorophore conjugates were diluted in PBS and stored at 4°C. Antibody concentration and degree-of-labeling (DOL) were determined with Nanodrop.

### Immunostaining and Ce3D clearing of tissue slices

Tissue slices were incubated in Mouse BD Fc Block (1:100; BD Biosciences) prepared in 500µl BD Perm/Wash buffer (1:10 diluted in distilled H_2_O; BD Biosciences) for 24 hours at room temperature (RT). For primary antibody staining, blocking buffer was replaced with antibody cocktail prepared in 400µl Perm/Wash buffer containing a mixture of fluorophore-conjugated antibodies (1:50 to 1:100 dilution) and incubated for 72 hours at RT on a slow shaker (60-70 rpm). Stained tissues were then washed in 2ml Perm/Wash buffer for another 24 hours. Following the wash step, post-fixation of the tissue was performed by first replacing the Perm/Wash buffer with 2ml PBS for 30 minutes, followed by post-fixing in 500µl 1% PFA for 15 minutes at RT, and then washed again with 2ml PBS for 30 minutes. Post-fixed tissues were then transferred into Ce3D solution (see below) for clearing. All steps were performed in the dark in 24-well plates covered in aluminum foil to protect tissues and fluorophores from light exposure.

For samples with secondary antibody staining, following the wash step after primary staining, the Perm/Wash buffer was replaced with 500µl Perm/Wash buffer containing fluorophore-conjugated secondary antibodies (1:500 dilution) and incubated for another 48 hours, washed for 24 hours and then followed by post-fixation step as described above. Antibodies used are listed in **Supplementary Table 1**.

For tissue clearing, Ce3D tissue clearing solution was prepared as described previously^12^ with modifications. Briefly, a 10ml clearing solution was prepared using 5.5ml 40% (v/v) N-methylacetamine (diluted with PBS; Sigma-Aldrich) and 10g Histodenz (Sigma-Aldrich) without Triton X-100 detergent. The tube containing Ce3D solution was then incubated in a heated shaker at 37°C and 150 rpm for at least 1 hour until the solid powder was fully dissolved. The prepared Ce3D solution was then stored on a slow shaker at RT, wrapped in aluminum foil to protect from light exposure. The refractive index of Ce3D was measured using a digital refractometer, with the expected value of ∼1.52.

For tissue clearing, small chambers containing ∼700µl Ce3D solution were prepared on a glass slide (SuperFrost Plus, VWR) by stacking two CoverWell incubation chambers (0.5mm depth with a 13mm chamber diameter; Grace Biolabs) on top of an adhesive SecureSeal imaging spacer (Grace Biolabs) to prevent leakage between the silicone chamber and the glass slide. Post-fixed tissue slices were transferred into Ce3D-filled chambers and sealed with a glass coverslip. Tissues were cleared for at least 48-72 hours on a gentle shaker (60-70 rpm) at RT prior to imaging.

Before imaging, a shallow imaging chamber was created by stacking 2 layers of adhesive SecureSeal imaging spacers (2 x 0.13mm depth) on a SuperFrost Plus glass slide. Up to 4 LN tissue slices were placed into an imaging chamber of 20mm x 20mm (spacers of 20mm circular chamber diameter trimmed to a square using a scalpel blade), arranged in a 2×2 grid, for batch imaging of tissues from the same experiment. The shallow chamber was then filled with Ce3D solution and gently sealed with a No. 1.5 glass coverslip (VWR).

### Laser Scanning Confocal Microscopy

Volumetric images were acquired using an inverted Leica Stellaris or upright TCS SP8 X confocal microscopes (Leica) equipped with a pulsed white light laser and four tunable spectral Hybrid Detectors (HyD). Images were acquired with a 20x multi-immersion objective (with correction collar adjusted to oil immersion), NA = 0.75 and working distance of 0.66mm and captured at a digital zoom of 1.5 (0.361µm xy pixel resolution) and 2µm z-step over the full thickness of the tissue slice. Excitation of CellTrace Violet and eFluor 450 was performed using a fixed 405nm laser line. 12- and 16-bit images were typically acquired, although some experimental datasets were acquired as 8-bit images. Image tiles were taken with 5% overlap and merged using the Leica LAS X Navigator application.

For high-resolution imaging of protein co-localization, a digital zoom factor of 2.0-2.5 (0.227-0.284µm xy pixel resolution) and 1.5µm z-step were used. For anti-NFAT1 stained tissues, images were acquired with a digital zoom factor of 2.0 (0.284µm xy pixel resolution) with 2µm z-step to provide sufficient lateral resolution for distinguishing cytoplasmic and nuclear NFAT localization.

### Chemical bleaching of fluorophores

For imaging experiments involving IBEX iterative staining^64^, imaged cleared tissue slices were first returned to wells filled with PBS for removal of clearing reagent until the tissue appearance became opaque. Tissues were then transferred into 2ml Perm/Wash buffer and washed for 24 hours at RT, then transferred into new Perm/Wash buffer-containing wells for two subsequent washes (24 hours each) to minimize retention of clearing reagent within the tissues.

To chemically bleach fluorophores with lithium borohydride (LiBH_4_; STREM Chemicals, cat no. 93-0397), bleaching solution was prepared at 10mg/ml LiBH_4_ concentration in distilled water, as described previously^64^. Tissue slices were then washed in PBS for 30 min at RT, replaced with LiBH_4_ solution for 45 min and placed ∼30 cm from a fluorescent light source. Bleached tissues were then washed again in PBS for 30 min, and finally transferred into a new well containing antibody cocktail solution for subsequent staining steps.

### Image Analysis

#### Image pre-processing, segmentation and histo-cytometric quantification of single cells

Please refer the **Supplementary Methods** for detailed steps of the image processing and analysis pipeline. Briefly, a Python-based computational pipeline was developed and optimized to enable distributed processing of large volumetric datasets on the NIH HPC Biowulf cluster. Raw image data were pre-processed to compensate for spectral spillover and correction for intensity attenuation along the z-axis.

Single cell segmentation was performed with a modified version of Stardist3D^65^. A small image region was cropped from a representative image and manually annotated as training dataset, and the custom trained model was used for segmentation of all image datasets from the same experiment. In most cases, Ki-67 nuclear stains were used as the segmentation channel. The nuclear masks were then morphologically dilated to encompass membrane/cytoplasmic region of the cells, and subtraction of both masks generated a new membrane/cytoplasmic mask. Mean intensity for each channel was determined by summing the masked voxel intensities divided by the sum of all mask voxels for every cell. Output arrays containing both cell coordinates (x, y, z) and mean marker intensities were exported for downstream analyses.

For histo-cytometry gating of single cells, custom Python-based scripts were utilized to visualize marker intensities as two-dimensional histo-cytometric plots and for gating on further subsets. Donor OT-I cells were selected based on Ki-67 and GFP/CD45.1. For gating polyclonal activated CD8+ T cells, Ki-67 and CD8b were used. Further subsets were generated based on TCF-1 and PD-1 expression. The spatial distribution of each subset was then visualized using Imaris 10.0 (Bitplane).

Quantification of marker expression for each subset was visualized with violin plots using the Python-based *seaborn* visualization library. PCA analysis was performed using Python-based *scikit-learn* library on z-score-normalized marker intensity and plotted using *matplotlib* library to visualize relative marker expression.

For quantification of TCF-1 expression level among OT-I cells in close proximity to dense cDC1 region, the threshold for cDC1-dense^hi^ gate was determined using the mean + 1 standard deviation of cDC1 density normalized to the max cDC1 density value of the OT-I population (see ‘cDC1 density’ section). OT-I cells above the cDC1-dense threshold (mean + 1 s.d.) were then gated (cDC1-dense^hi^) and their relative TCF-1 expressions (normalized to naïve T cell expression) displayed as violin plots. Naïve TCF-1 expression was determined by selecting a small patch (∼512×512) of the LN T cell zone densely populated with naïve T cells expressing high level of TCF-1 and calculating the mean intensity of the TCF-1 channel. Further gating on TCF-1^hi^ subset was performed by selecting cells with TCF-1 value (normalized to naïve T cells) of >0.7.

#### cDC1 density

For determining spatial proximity to dense cDC1 region, a Gaussian smoothing filter of bandwidth σ=3.6µm (10 pixels) was applied to the XCR1 channel for each z slice. The mean intensity obtained from a cell’s nuclear mask yields the spatial density of cDC1 at the cell’s centroid.

#### Kernel density estimation

3D image coordinates of cells of interest were converted to world coordinates by multiplying their voxel dimensions. Kernel density estimation was then performed using the *TreeKDE* module from *KDEpy* library, with an isotropic Gaussian kernel (bandwidth σ=6µm) across a grid system of 10µm interval in each axis. Weights were set to normalized protein marker expression (e.g., PD-1, SLAMF6) of each cell. To generate a kernel density map for visualization, the density values were summed over a selected z-axis range comprising 80µm thickness (out of ∼300µm full volume thickness) to reduce clutter. A contour map was then generated using the *contourf* function in *matplotlib* library set to a perceptually uniform colormap for visualization of the probability density on a two-dimensional plot.

#### Heatmap visualization

Each parameter (protein expression, spatial density) was first standardized to obtain a z-score for each cell. The cell population was then sub-divided and sorted based on the manual gating strategy defining SLAMF6+ T_SL_, SLAMF6-T_SL_ and TCF-1-T_EFF_. A perceptually uniform colormap was applied to display the relative z-score of each parameter. Note that both screen and print resolutions are not sufficient to enable discrimination of single cells within the heatmap, which contains the total population of OT-I recovered from the LN tissue slice (>65k cells in the dataset shown in Fig. 1e).

#### Visualization

For visualization of protein markers expressed by gated cells, segmented labels (nuclear masks) of the gated cells were processed with the morphological dilation tool (radius=6) from *scikit-image* library to generate a new mask encompassing the cytoplasmic/membrane region of the cells. Individual image channels were then multiplied with the mask to generate a new image displaying only protein marker intensities masked within the cells of interest. These masked images were then visualized using Imaris 10.0 (Bitplane). To visualize the relative protein expression level, perceptually uniform colormaps were used and the min/max scaling (contrast and brightness) of each channel individually adjusted to avoid under- and over-saturation of the marker intensity.

#### Animation

For animation of 3D imaging dataset, the *napari-animation* library was used and a custom animation script was made to generate keyframes specifying the camera positions and angles, image layers’ colormap, adjustments of contrast/brightness, as well as clipping planes for animating transition between layers and for focusing on thin cross-sections of the imaging volume. The output movie file was further processed and edited in Davinci Resolve 18.6 (Blackmagic Design) to include annotations with text and graphic items.

### Tissue preparation for flow cytometry

Lymphocytes were isolated from lymph nodes and spleens and made into single-cell suspensions using a syringe plunger and 100µm or 70µm cell strainers (MACS SmartStrainer, Miltenyi Biotec). ∼3×10^6^ cells were used in subsequent staining steps for flow cytometry analysis. Dendritic cell isolation was performed as described previously^66^. Briefly, lymphoid tissues were sliced into small fragments using a scalpel blade and incubated in a digestion mix of collagenase type III (Worthington, 1mg/ml) and DNase I (20ug/ml) and vigorously mixed for 25 min at room temperature, followed by addition of 0.1M EDTA solution at 1/10 digestion volume for 5 min to dissociate lymphocytes from dendritic cells. Tissue debris were filtered out by passing the cell suspension through a 70µm nylon mesh.

T cells from skin tumor samples were isolated as described previously^67^. Briefly, a 1cm^2^x1cm^2^ tumor-containing skin patch was harvested into collagenase type III (Worthington, 3mg/ml), finely chopped with scissors and incubated at 37°C for 90 min before pressing through 70µm cell strainers. For spleen and tumor samples, cells were also treated with red blood cell lysis buffer prior to staining. Cell counts in LN and spleen were determined using an automated Cellometer T4 cell counter (Nexcelom Bioscience).

### Flow Cytometry

For detection of polyclonal OVA-specific CD8+ T cells, cells were first incubated with PE-conjugated H-2K^b^-SIINFEKL tetramer (1:100, NIH Tetramer Core) for 20 min at 37°C, washed, followed by cell surface marker staining for 25 min at 4°C. Mouse BD Fc Block (1:200; BD Biosciences) was also included during cell surface marker staining step. A fixable LIVE/DEAD near-infrared staining dye was used for determining cell viability. For detection of intracellular proteins, stained cells were further treated with fixative and stained for antibodies against intracellular proteins using Foxp3/transcription factor staining buffer kit per the manufacturer’s protocol (eBioscience). For tumor samples, CountBright Plus Absolute beads (Thermofisher) were added prior to sample acquisition. Samples were acquired using a BD Fortessa (BD Biosciences) and analyzed using Flowjo 10 (Treestar). Antibodies used are listed in **Supplementary Table 1**.

### Affinity index

To estimate the TCR affinity of tetramer-stained cells, an affinity index was derived from dividing tetramer staining intensity by TCR (CD3) staining intensity (tetramer/CD3 ratio). This is possible because tetramers elute from labeled T cells in the wash buffer during incubation steps in reasonable proportion of affinity, in particular the off-rate of the interaction. When normalized to expressed TCR level, we can use the absolute staining as a proxy for this off-rate and hence, affinity. Cells were post-fixed after the surface marker staining step, prior to intracellular antibody staining, to minimize elution of tetramers over time. Given that tetramer binding varies from batch-to-batch and is more sensitive to incubation conditions compared to antibody staining, a z-score for each subset from each animal was derived from the log-affinity index (log tetramer/CD3 ratio) of the tetramer-stained cells, standardized with reference to SLAMF6+ T_SL_ of the control IgG-treated group from the same experimental cohort. The mean of the z-score, Mean z(Log(Affinity Idx)), thus estimates the distribution of TCR affinity of each subset and treatment group in relation to SLAMF6+ T_SL_ of the control IgG-treated group.

A normalized ratio was also derived from the TCR affinity indices of tetramer-stained cells, by normalizing the affinity indices of each subset from each animal to the reference affinity index of SLAMF6+ T_SL_ from control IgG-treated group pooled from the same experimental cohort. This normalized ratio, Mean Affinity Idx, estimates the relative TCR binding affinity of each subset and treatment group in relation to SLAMF6+ T_SL_ of the control IgG-treated group. When specified, other reference subsets (e.g., Day 6 TCF-1+ T_SL_) were also used for normalization.

### Modeling of TCR Affinity and Cell States

In the course of these studies, we observed a number of outlier responses among the treated individual mice. Such results are not unexpected based on the known variation in TCR repertoire between inbred animals^13^. A linear modeling analysis was conducted to determine if these outlier events affected the conclusions of our flow cytometry studies.

The TCR affinity of single cells was quantified as the log-ratio of tetramer staining to CD3 staining (log tetramer/CD3 ratio). Within each experiment, these log-ratios were standardized with reference to the control IgG-treated SLAMF6+ T_SL_, to emphasize the differences in affinity between cell states and treatment groups. The three major subsets of OVA-tetramer+ CD8+ T cells: SLAMF6+ TCF-1+ T_SL_, SLAMF6-TCF-1+ T_SL_, and TCF-1-T_EFF_ were defined as distinct cell states. Standardization was performed separately for each experiment, to control for experiment-to-experiment staining variability, because affinity measurements differed substantially across experiments. Cellular affinity was then regressed on cell state, treatment, and their interaction using a Bayesian multilevel model with a Gaussian likelihood. Hyperparameters for the effect of mouse identity on the intercept and on the cell state slopes were included to account for mouse-to-mouse variability. Predictions from this model were simulated without mouse-to-mouse variability to isolate the effects of cell state and treatment on the expected distributions of cellular affinity.

In a second model, the number of cells in each cell state from each animal were log-transformed. These log-counts were then standardized across all groups of cells. Cell count was regressed on cell state, treatment, and their interaction using a Bayesian multilevel model with a Gaussian likelihood. Hyperparameters for the effect of mouse identity on the intercept and on cell state slopes were included to account for mouse-to-mouse variability. Predictions from this model were simulated without mouse-to-mouse variability to isolate the effects of cell state and treatment on the expected cell counts.

The simulated predictions from the first model provided expected distributions of log-affinity relative to the mean of control IgG-treated SLAMF6+ T_SL_ for each combination of cell state and treatment. The fraction of cells with log-affinity greater than the mean of control IgG-treated SLAMF6+ T_SL_ was simply the fraction of cells above 0 in each distribution. These fractions were multiplied by the expected total number of cells for each combination of cell state and treatment, provided by the simulated predictions of the second model. This gave the expected number of cells greater than the mean log-affinity of control IgG-treated SLAMF6+ T_SL_ (“above average TCR affinity threshold”) for each combination of cell state and treatment.

### Statistics

Statistical tests were performed in Prism 9.0 software (GraphPad). Data analyses were performed with unpaired, two-tailed Student’s *t*-test, or one-way ANOVA with Tukey’s post-hoc multiple comparison tests, as specified in the text or figure legends. p<0.05 was considered significant: *p<0.05, **p<0.01, ***p<0.001, ****p<0.0001.

## Supplementary Methods

Supplementary Methods.pdf

Detailed descriptions of image analysis methods.

## List of Supplementary Table

**Supplementary Table 1**. Supplementary_Table_1.xlsx

Excel file containing list of antibodies used in this study.

## List of Supplementary Videos

**Supplementary Video 1.** Supplementary_Video_1.mp4

Quantitative 3D tissue microscopy pipeline for mouse tumor-draining lymph node.

**Supplementary Video 2.** Supplementary_Video_2.mp4

Histo-cytometric analysis of antigen-specific CD8+ T cells in the tumor-draining lymph node.

**Supplementary Video 3.** Supplementary_Video_3.mp4

Single-cell level phenotyping of antigen-specific T cells.

---

**Extended Data Figure 1.**
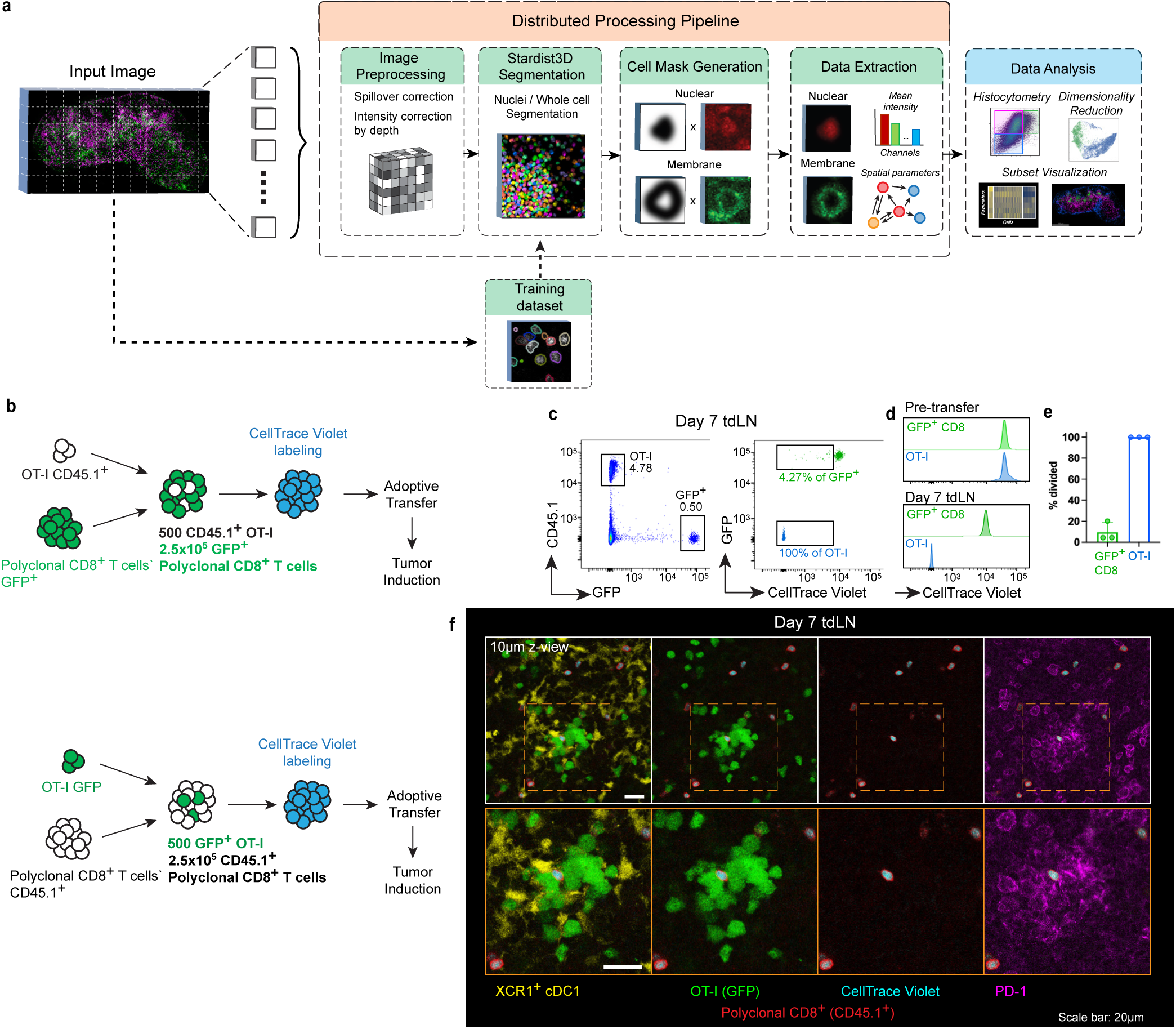
Imaging data processing pipeline and tracking of antigen-specifiic CD8+ T cell proliferation in the tdLN. **a**, Schematics of a distributed image processing and analysis pipeline that enables fast and accurate single-cell level analysis of large 3D volumetric datasets. The raw input image is divided into sub-blocks, which are processed piece-wise in parallel to perform image corrections, cell segmentation, masks generation and data extraction. The training dataset for segmentation is generated from semi-automated label annotation from a small, cropped region of a representative image from the same experiment. Further downstream analyses are performed with extracted data. **b**, Experimental scheme of co-labeling of OT-I and polyclonal CD8+ T cells isolated from naïve mice with CellTrace Violet proliferation dye, in reciprocal formats: OT-I.CD45.1 and CD8.GFP (top: **c-e**); OT-I.GFP and CD8.CD45.1 (bottom: **f**). Cells were co-transferred into recipient mice at 1:500 ratio. **c**, Left, flow cytometry plot of donor cell populations recovered from Day 7 tdLN and right, their corresponding CellTrace Violet dye intensity. Gates denote populations with diluted CellTrace Violet signal (CTV^low^). **d**, Histograms showing CellTrace Violet signal measured from pre-transfer cell suspension (top) and those recovered from Day 7 tdLN (bottom). **e**, Quantification of proportion of divided cells (CTV^low^) from Day 7 tdLN as gated in (**c**). **f**, Representative cross-section images showing clustering PD-1+ OT-I.GFP cells (green) and polyclonal non-antigen specific CD8+ T cells (CD45.1, red) in Day 7 tdLN. Proliferation dye (cyan) has been fully diluted among the antigen-specific OT-I.GFP population. Dashed box indicates the area shown in the close-up images (bottom).

**Extended Data Figure 2.**
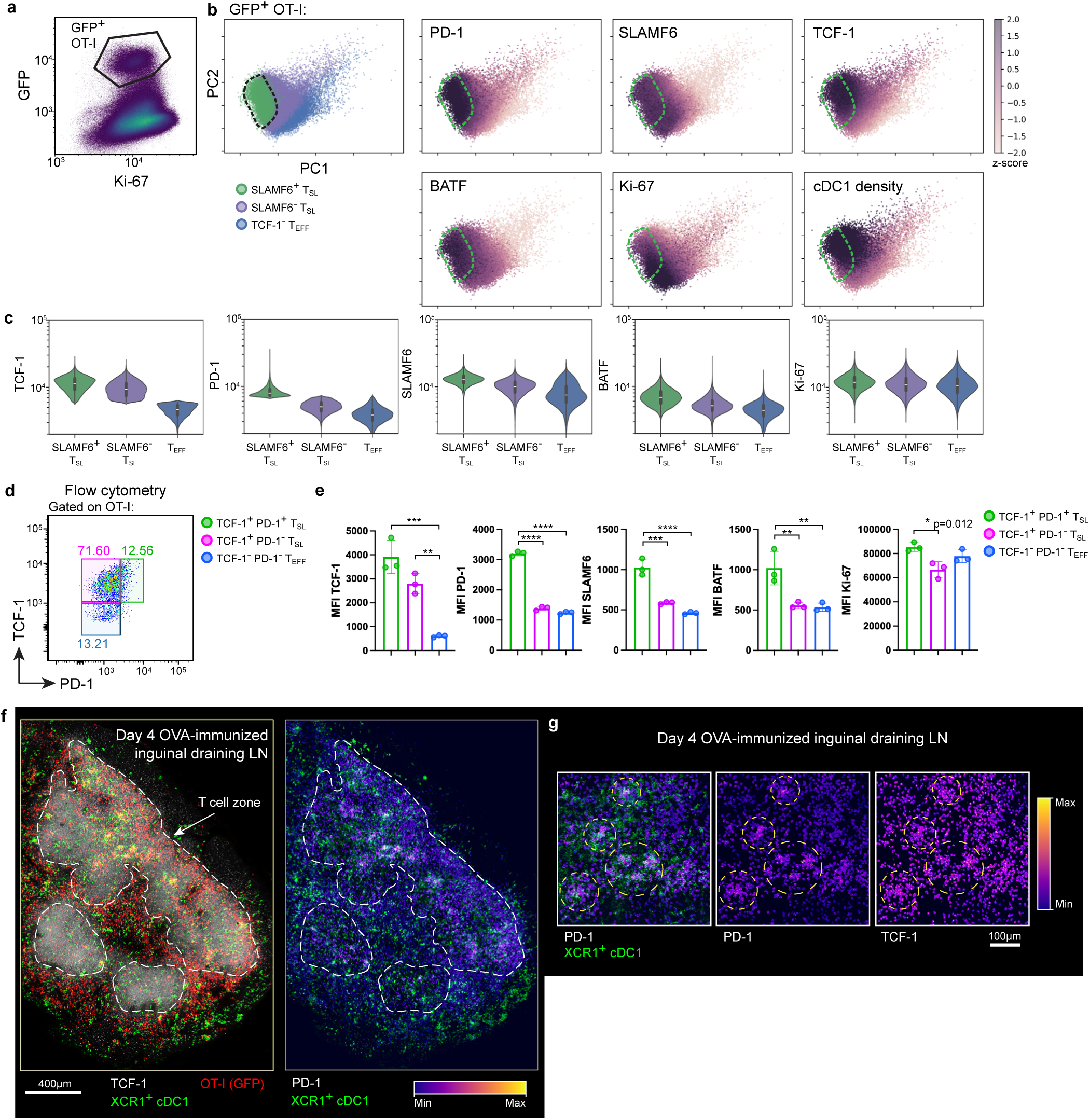
Phenotypic and functional state analysis of tumor antigen-specific OT-I cells in Day 8 tdLN. **a**, Representative histo-cytometric plot showing gating strategy of OT-I cells in Day 8 tdLN based on GFP and Ki-67 co-expression. **b**, PCA plots of color-coded OT-I subsets based on TCF-1 and PD-1 expression (Fig. 1c) (left panel) and the standardized expression for each marker (right panels). Dotted region indicates TCF-1+ PD-1+ SLAMF6+ T_SL_. **c**, Protein expression levels for each OT-I subset as defined in Fig. 1c. Data shown are representative of 2 independent experiments (n=2-3 per experiment) **d**, Flow cytometric plot of OT-I cells recovered from Day 7 tdLN with subsets gated similarly to histo-cytometric data from Fig. 1c. **e**, Protein expression levels for each OT-I subset as gated in (**d**). **f-g**, Day 4 inguinal dLN from mice transferred with 500 OT-I.GFP and immunized with OVA/poly(I:C) on the flank skin. **f**, Cross-section image showing the demarcation of T cell zone (dashed boundaries) based on TCF-1 staining (left) and PD-1 intensity (masked on OT-I.GFP cells) (right). **g**, Close-up images showing co-localization of PD-1+ OT-I cells with cDC1, PD-1 intensity only and TCF-1 intensity only (both masked on OT-I cells) in the order from left to right. Data are representative of 2 independent experiments (n=2 per experiment). Error bars: means ± s.d. *p<0.05, **p<0.01, ***p<0.001, ****p<0.0001; one-way ANOVA with Tukey’s multiple comparisons.

**Extended Data Figure 3.**
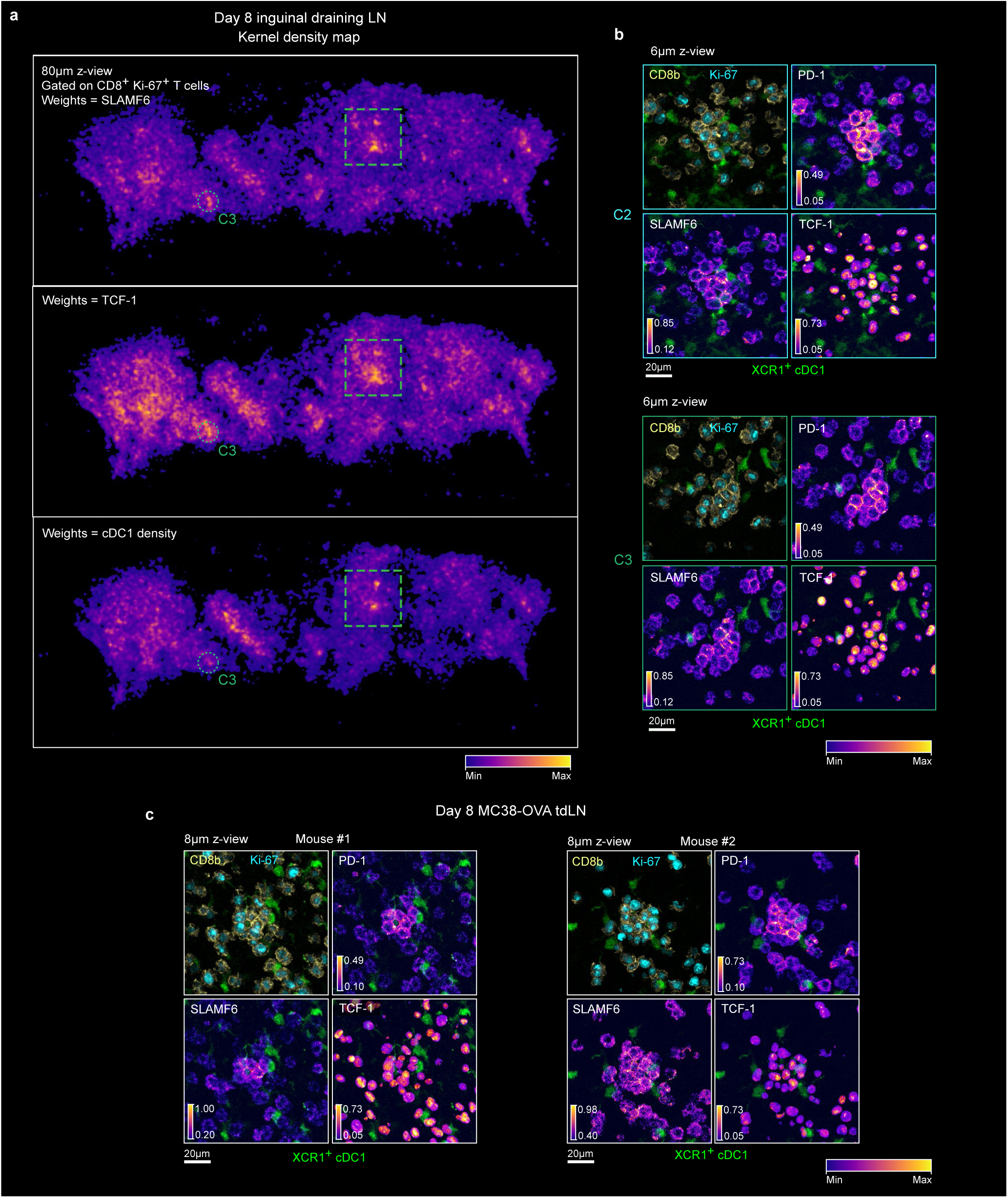
Distinct foci of polyclonal CD8+ SLAMF6+ T_SL_ in Day 8 tdLN. **a**, Kernel density map of activated polyclonal CD8+ T cells in Day 8 tdLN. Density was estimated using a Gaussian kernel of 6µm bandwidth, weighted on SLAMF6 (top), TCF-1 (middle) or cDC1 density (bottom). Green dashed box indicates close-up region corresponding to Fig. 2b-c, while dotted circle (C3)

**Extended Data Figure 4.**
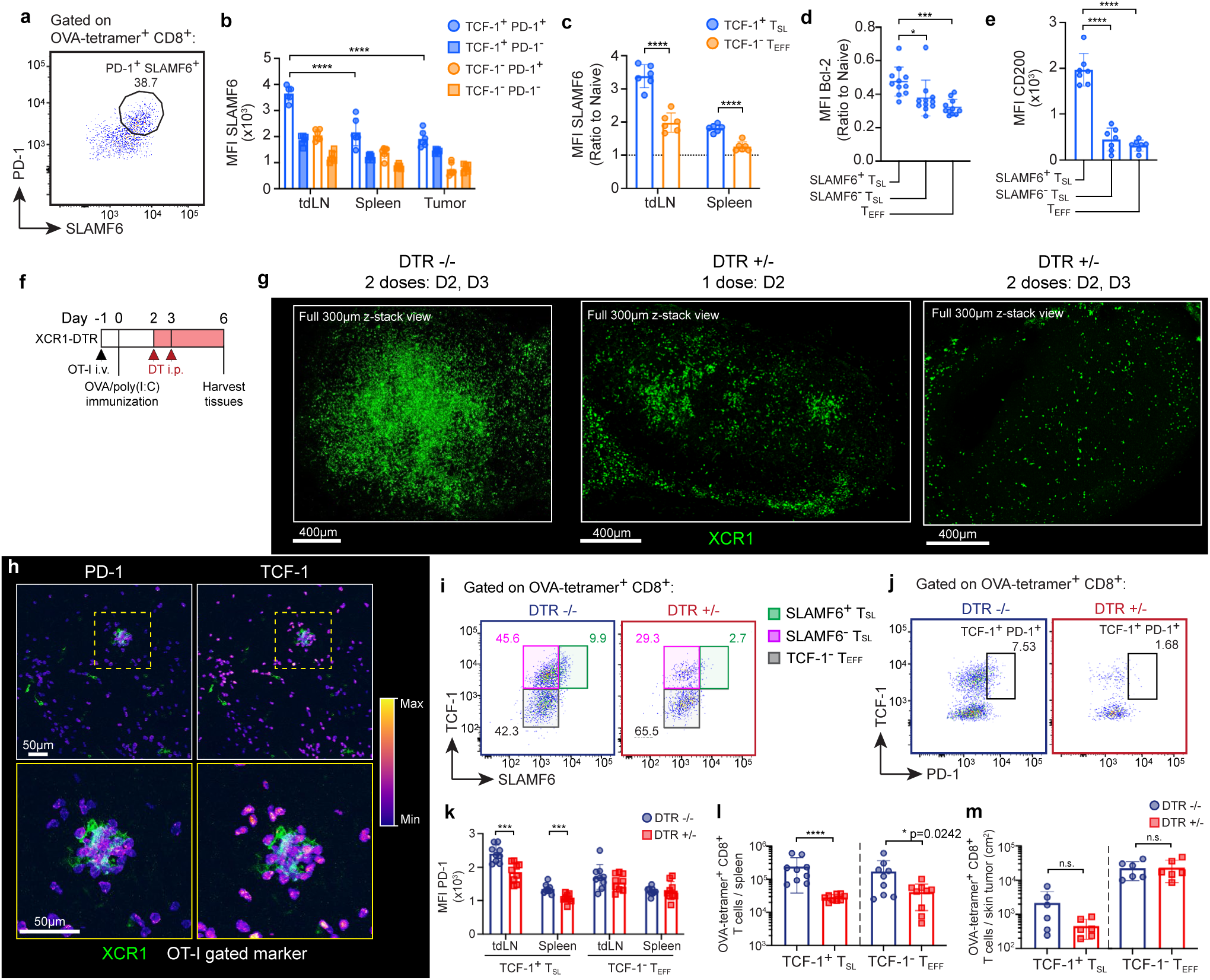
Characterization of SLAMF6+ T_SL_ across tissues and late antigen presentation niche driven expansion of T_SL_. **a**, Flow cytometry plot showing correlation of PD-1 and SLAMF6 protein expression on OVA-tetramer+ CD8+ T cells from Day 10 tdLN. **b**, Mean SLAMF6 fluorescence intensity on different subsets of OVA-tetramer+ cells across different tissues. SLAMF6 expression was particularly enriched in TCF-1+ T_SL_ residing in tdLN, as shown by the normalized SLAMF6 expression (normalized to naïve TCF-1+ PD-1-CD8+ T cells) in (**c**). **d**, Bcl-2 expression (normalized to naïve TCF-1+ PD-1-CD8+ T cells) and **e**, mean fluorescence intensity of CD200 expressed on each subset of OVA-tetramer+ CD8+ T cells. Data from 2 independent experiments (n=6), and 3 independent experiments (n=11) for (**d**). **f**, Experimental scheme of diphtheria toxin (DT) administration in mice transferred with OT-I cells and received intradermal OVA/poly(I:C) immunization. **g**, Full 300µm-thickness view of Day 6 immunized inguinal dLN stained with polyclonal anti-XCR1 antibody (green) to assess depletion efficiency. **h**, Close up images of OT-I clustering around one of the few remaining cDC1, showing PD-1 (left) and TCF-1 (right) expression masked on OT-I. Dotted yellow box denotes a close-up region shown in the bottom row. **i,** Gating strategy to quantify SLAMF6^+^ T_SL_ subset in Fig. 3d. **j-k**, Representative flow cytometric plots showing the loss of PD-1+ OVA-tetramer+ CD8+ T cells after DT depletion of cDC1 (**i**) and quantification of mean PD-1 intensity (**j**) in Day

**Extended Data Figure 5.**
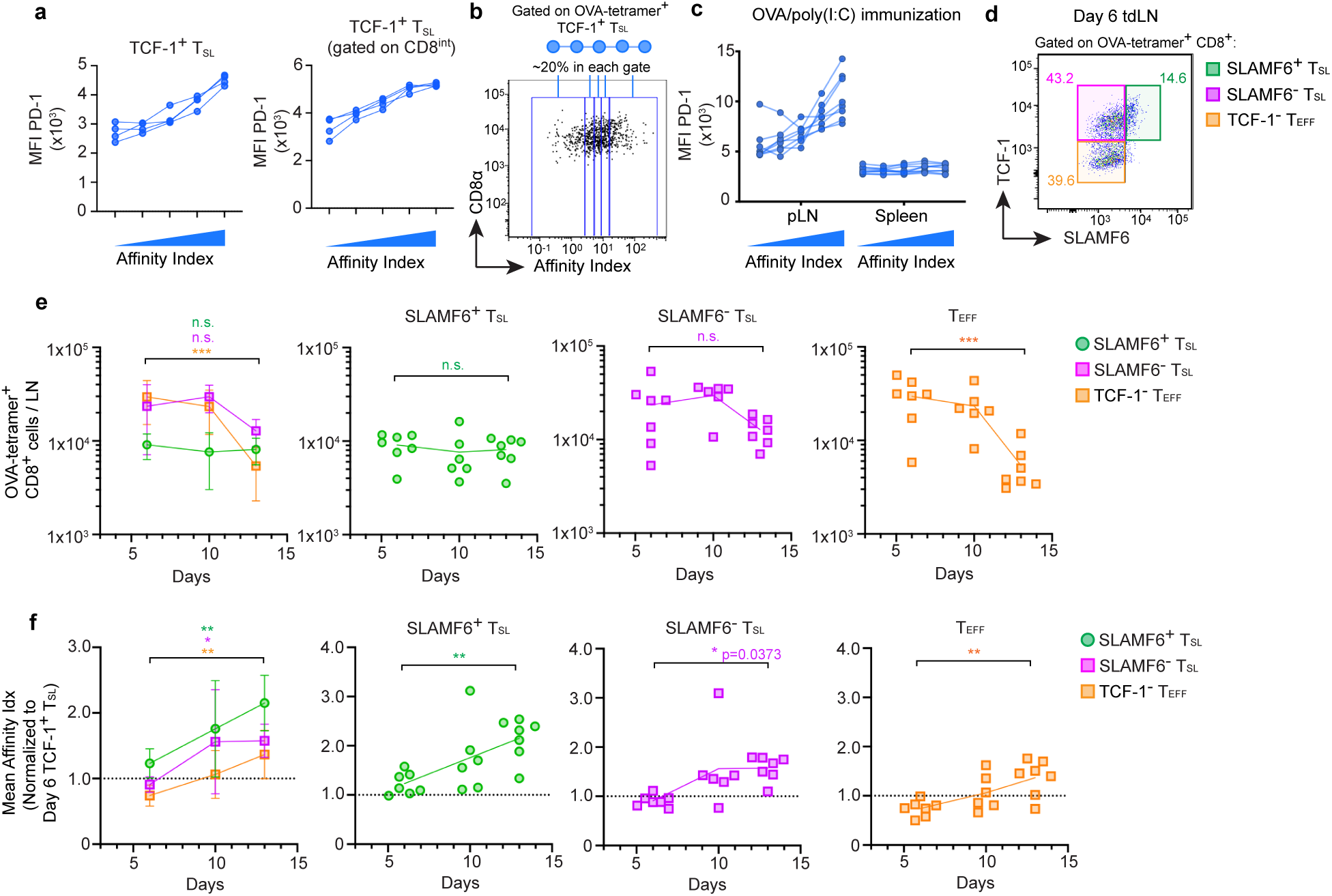
PD-1 expression of T_SL_ correlates with TCR affinity and drives affinity evolution in the draining LN. **a**, Mean fluorescence intensity of PD-1 expression on OVA-tetramer+ TCF-1+ T_SL_ across different bins of affinity indices (as defined in Fig. 3e), gated on the entire sub-population (left panel) or along a band of cells with intermediate CD8α expression (∼50% of the sub-population, right panel). **b**, Flow cytometry plot showing gating strategy of binning OVA-tetramer+ TCF-1+ T_SL_ in Day 5 draining popliteal LN of OVA immunized mice, based on TCR affinity indices. Mean fluorescence intensity of PD-1 in the dLNs and spleens is shown in (**c**). Data pooled from 2 independent experiments (n=8). **d**, Gating strategy for OVA-tetramer+ CD8+ T cell subsets in (**e, f**). The absolute count of each subset (**e**) and the average TCR affinity indices (normalized to Day 6 TCF-1+ T_SL_) (**f**) shown combined (left) or as individual subsets. Data from 2 independent experiments (n=6-8 per time point). Error bars: means ± s.d. *p<0.05, **p<0.01, ***p<0.001; one-way ANOVA with Tukey’s multiple comparisons.

**Extended Data Figure 6.**
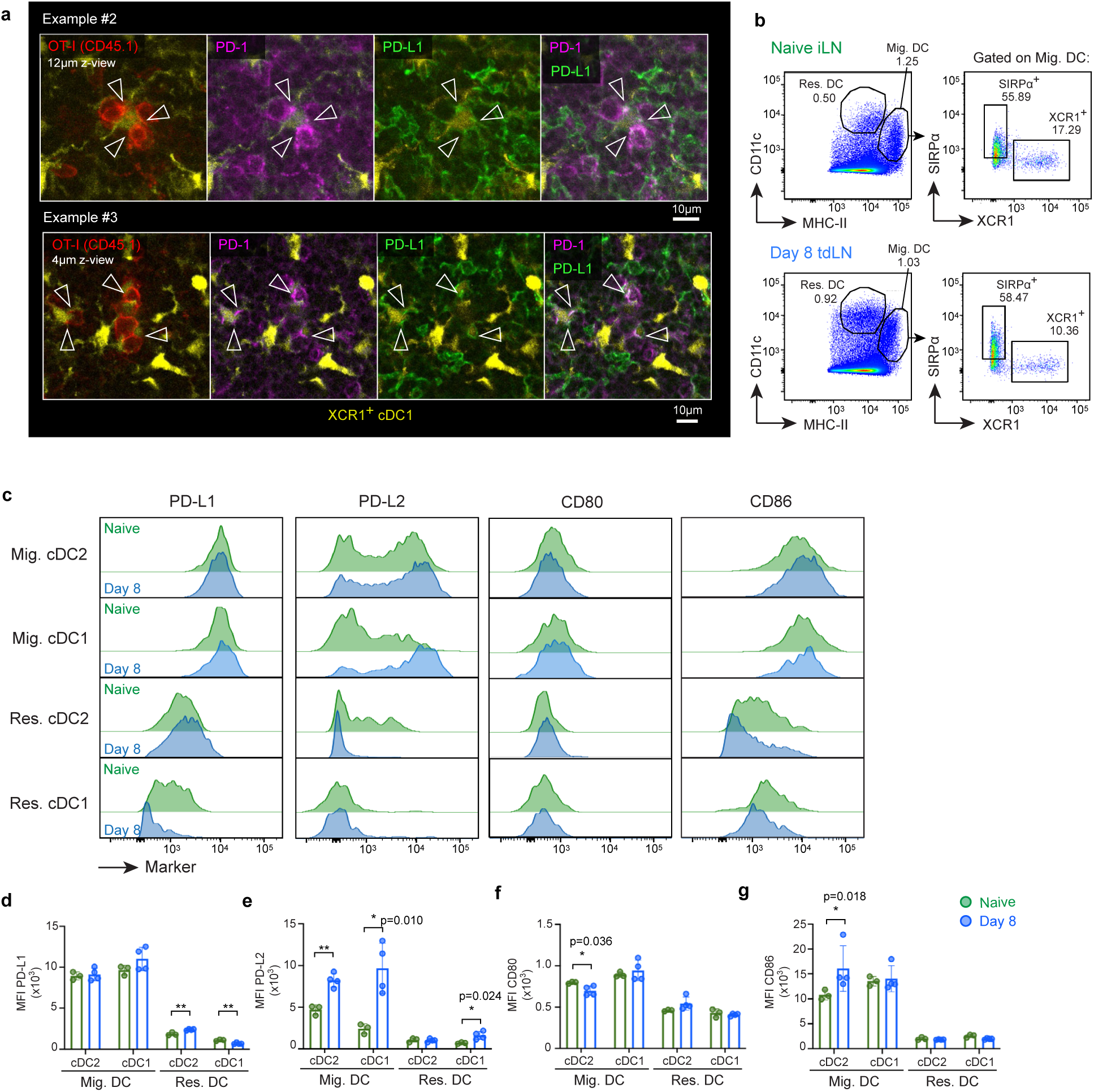
PD-1/PD-L1 engagement in late antigen-presenting T-DC niches. **a**, Multi-panel images showing thin cross-sections (12µm and 4µm z-thickness) of two examples of clustering OT-I cells and the co-localization of PD-1 (magenta) on OT-I.CD45.1 (red) with PD-L1 (green) expressed on cDC1 (yellow), as in Fig. 4b. Arrows indicate polarization and punctate microcluster formation of PD-1 and their co-localization with PD-L1 blending into white pixels. Data representative of at least 2 independent experiments (n=2-3 per experiment). **b**, Flow cytometric gating strategy of migratory versus LN-resident dendritic cell subsets isolated from naïve (top) or Day 8 tdLN (bottom) based on CD11c and MHC-II expression (left panel), and further into SIRPα+ cDC2 and XCR1+ cDC1 (right panel). Histogram panels showing relative expression of PD-L1, PD-L2, CD80 and CD86 markers on each cDC subset are shown in (**c**) and the quantification of mean fluorescence intensity for each marker in (**d**). Data from 2

**Extended Data Figure 7.**
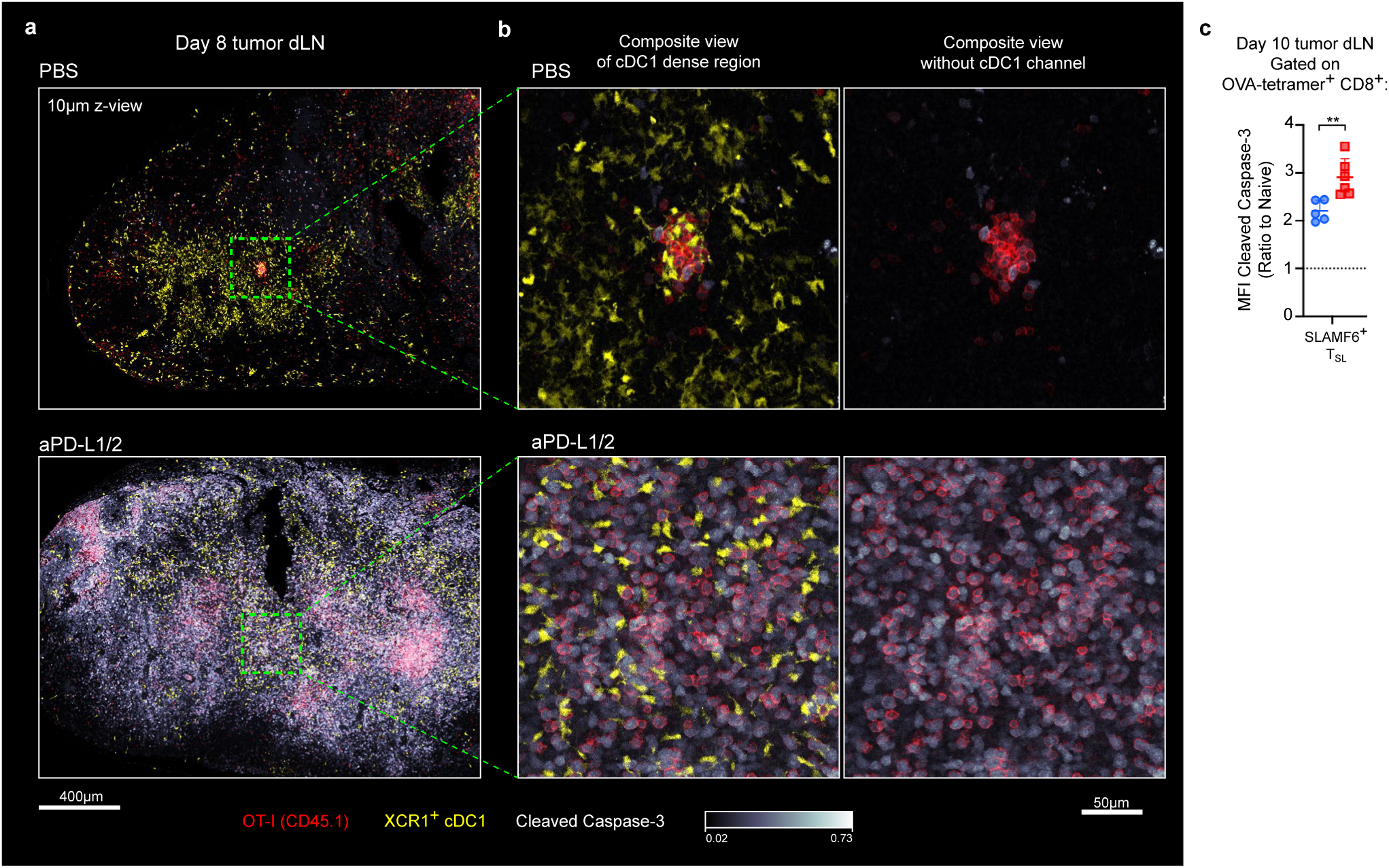
PD-1 checkpoint blockade induces apoptosis of CD8+ T cells in the tdLN. **a**, Cross-section of Day 8 tdLN from mice treated with PBS or anti-PD-L1/PD-L2 blocking antibodies following the experimental scheme from Fig. 4c. Cleaved caspase-3 staining indicates increased apoptosis of the LN cells. Dotted box denotes the close-up of cDC1-dense region shown in (**b**), displayed with (left) or without cDC1 channel (right). Data are representative of 3 independent experiments (n=2-3 per group per experiment). **c**, Relative expression of cleaved caspase-3 of OVA-tetramer+ SLAMF6+ T_SL_ recovered from Day 10 tdLN treated with PBS (blue) or anti-PD-L1/2 (red) as measured by flow cytometry. Expression level is normalized to CD44-TCF-1+ PD-1-naïve CD8+ T cell population. Error bars: means ± s.d. **p<0.01; unpaired two-tailed *t*-test.

**Extended Data Figure 8.**
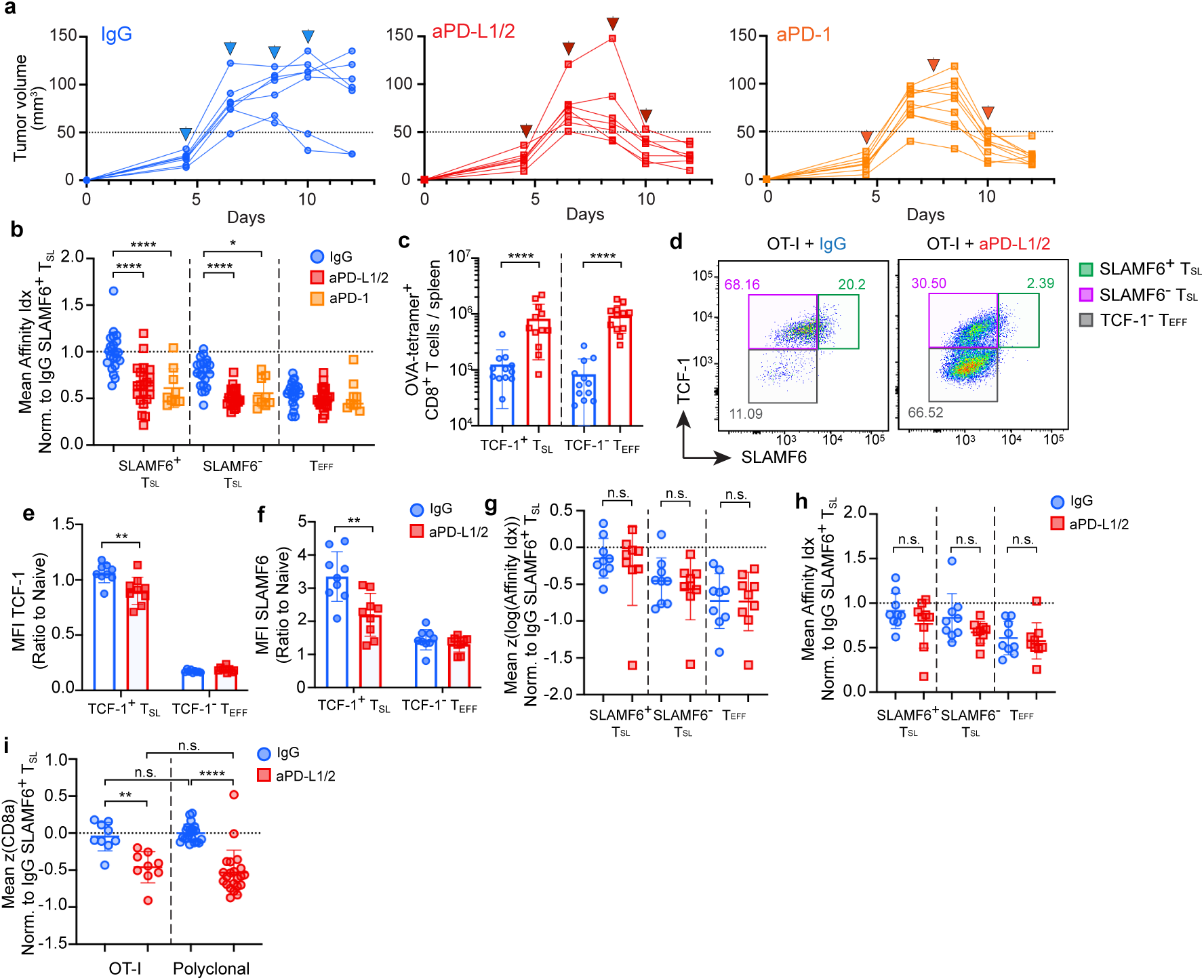
PD-1 checkpoint blockade reduces the frequency of high affinity CD8+ T_SL_ in the tdLN. **a**, Tumor growth curve in mice treated with control IgG, anti-PD-L1/PD-L2 or anti-PD-1. Arrows indicate checkpoint antibody injection. Data from 2 independent experiments (n=7 for IgG and anti-PD-L1/PD-L2; 9 for anti-PD-1). **b**, Mean affinity index of each subset of OVA-tetramer+ CD8+ T cells from Day 12 tdLN normalized to IgG-treated SLAMF6+ T_SL_ from the same experimental cohort. Data pooled from 7 independent experiments for IgG and anti-PD-L1/PD-L2 treatment (n=3-4 per group per experiment), 2 independent experiments for anti-PD-1 group (n=4-5 per experiment). **c**, Quantification of OVA-tetramer+ TCF-1+ T_SL_ and TCF-1-T_EFF_ recovered from Day 12 spleen. Data from 4 independent experiments (n=12 per treatment group). **d-i**, Tumor-bearing mice adoptively transferred with 2×10^3^ congenic CD45.1+ naïve OT-I.Rag1^-/-^ cells 1 day prior to tumor induction and treated with checkpoint antibody as shown in Fig. 5a. **d**, Representative flow cytometry plots showing changes in TCF-1 and SLAMF6 expression of monoclonal OT-I cells from Day 12 tdLN. Gates denote subsets used in (**g, h**). **e-f**, Mean fluorescence intensity of TCF-1 (**e**) and SLAMF6 (**f**) of OT-I cells normalized to endogenous naïve (CD44-TCF-1+ PD-1-) CD8+ T cells. **g-h**, Standardized log transformed TCR affinity index (**g**) and mean affinity index (**h**) of each treatment group, normalized to IgG-treated SLAMF6+ T_SL_ from the same experimental cohort. Each data point represents the mean subset value from each animal. **i**, Standardized CD8α expression of SLAMF6+ OT-I T_SL_ compared with polyclonal SLAMF6+ T_SL_ data from Fig. 5b-i, normalized to the mean CD8α expression

**Extended Data Figure 9.**
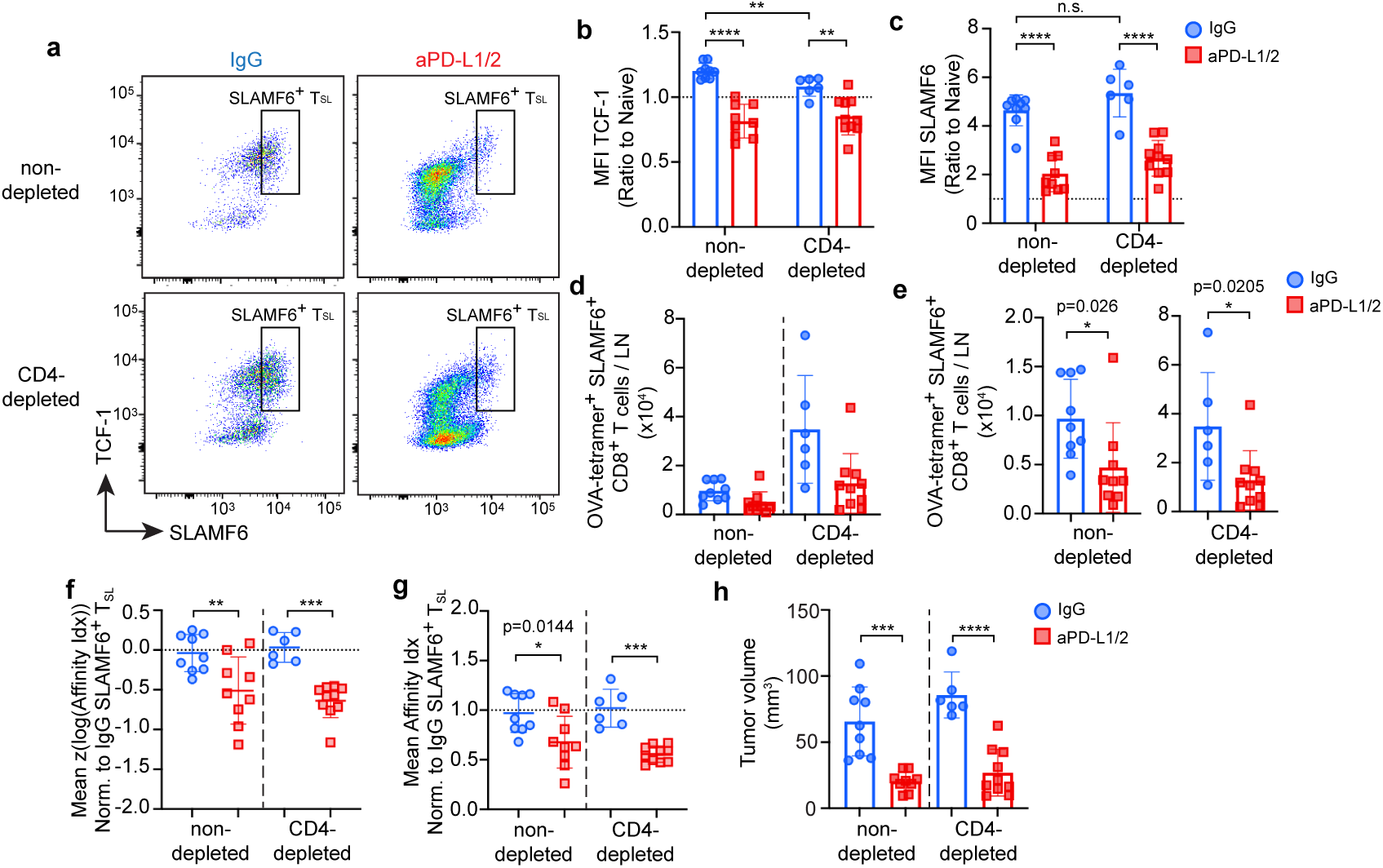
CD4+ T cells are not required to induce CD8+ phenotypic and clonal repertoire shift after checkpoint blockade. **a**, Representative flow cytometry plots of OVA-tetramer+ CD8+ T cells from Day 12 tdLN of mice pre-treated with or without anti-CD4 depleting antibodies, and with control IgG or anti-PD-L1/PD-L2 according to Fig. 5a experimental scheme. **b, c**, Mean fluorescence intensity of TCF-1 (**b**) and SLAMF6 (**c**) of OVA-tetramer+ TCF-1+ T_SL_ normalized to naïve (CD44-TCF-1+ PD-1-) CD8+ T cells. **d, e**, Quantification of OVA-tetramer+ SLAMF6+ T_SL_ from Day 12 tdLN (**d**), and re-scaled for each group (non-depleted and CD4-depleted) to show the relative extent of SLAMF6+ T_SL_ reduction after checkpoint treatment (**e**). **f, g**, Standardized log transformed TCR affinity index (**f**) and mean affinity index (**g**) of each treatment group, normalized to non-CD4-depleted, IgG-treated SLAMF6+ T_SL_ from the same experimental cohort. Each data point represents the mean subset value from each animal. **h**, Tumor volume measured at the experimental end point at Day 12. Data from 3 independent experiments (n=6-10 per treatment group). Error bars: means ± s.d. *p<0.05, **p<0.01, ***p<0.001, ****p<0.0001; unpaired two-tailed *t*-test in (**e**), one-way ANOVA with Tukey’s multiple comparisons in all others.

**Extended Data Figure 10.**
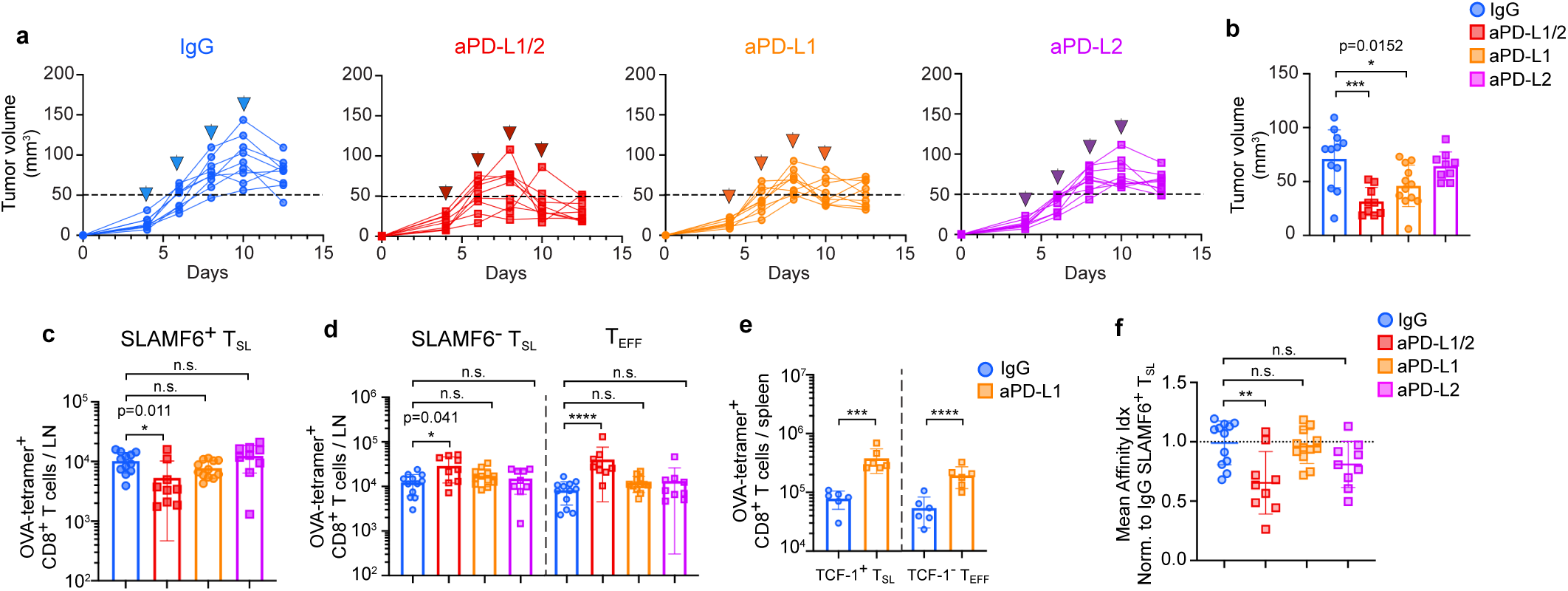
PD-L1 blockade alone does not result in dramatic loss of high affinity SLAMF6+ T_SL_. **a**, Tumor growth curve in mice treated with control IgG, anti-PD-L1/PD-L2, anti-PD-L1 only or anti-PD-L2 only. Arrows indicate checkpoint antibody injection. Tumor volume measured at experimental end point at Day 12 for each group is shown in (**b**). **c, d**, Number of OVA-tetramer+ SLAMF6+ T_SL_ (**d**), SLAMF6-T_SL_ and TCF-1-T_EFF_ (**e**) recovered from Day 12 tdLN. **e**, Number of OVA-tetramer+ TCF-1+ T_SL_ and TCF-1-T_EFF_ recovered from the spleen of mice treated with control IgG or anti-PD-L1 only at Day 12. **f**, Mean affinity index of each treatment group, normalized to IgG-treated SLAMF6+ T_SL_ from the same experimental cohort. Each data point represents the mean subset value from each animal. Data shown in (**a**) are from 3 independent experiments (n=9 per treatment group), (**b-d, f**) are pooled from 4 independent experiments for control IgG, anti-PD-L1 treatment groups (n=12), and 3 independent experiments for anti-PD-L1/PD-L2 and anti-PD-L2 groups (n=9). Data in (**e**) are from 2 independent experiments (n=6 per group). Error bars: means ± s.d. *p<0.05, **p<0.01, ***p<0.001, ****p<0.0001; unpaired two-tailed *t*-test in (**e**), one-way ANOVA with Tukey’s multiple comparisons in all others.

