## Supplementary Methods for "PD-1 controls differentiation, survival, and TCR affinity evolution of stem-like CD8+ T cells"

**Image Analysis**

Due to the very large number of cells acquired during 3D volumetric imaging (>1 million segmented cells), use of a conventional image processing pipeline conducted on dedicated workstations is impractical due to the exponential amount of data captured compared to thin tissue sections, with the length of time for processing increased by orders of magnitude (days-weeks).

To efficiently process such large 3D volumetric datasets, a Python-based computational pipeline was developed and optimized for distributed processing, in which the images were split into multiple sub-blocks to be processed in parallel on the NIH High Performing Computational (HPC) Biowulf cluster.

*Image Pre-processing*

Each raw Leica image file (.lif) was first converted into Imaris software format (.ims) using the Imaris File Converter tool (Bitplane). The ims format is a HDF5-based container and enables image processing steps to be seamlessly integrated into custom Python-based scripts.

The antibody staining panel was designed, together with the use of tunable excitation laser lines with tunable detection filters, to minimize spectral spillover and no further processing is required for most channels. For GFP/XCR1-venus signal crosstalk, pairwise compensation was performed as described previously[69]. Briefly, the spillover coefficient was first determined by sampling a small area containing GFP+ (fluorophore A) and venus+ (fluorophore B) cells, where the spillover coefficient is defined as:

$$S_{A>B} = \frac{A_{ch\_B}}{A_{ch\_A}}$$

where $S_{A>B}$ is the spillover coefficient for fluorophore A into channel B,

$A_{ch\_B}$ is the mean intensity of fluorophore A detected in channel B, and

$A_{ch\_A}$ is the mean intensity of fluorophore A detected in channel A as determined with a single-color control, or in cells with mutually exclusive fluorophores.

The reciprocal spillover coefficient for fluorophore B into channel A was also obtained:

$$S_{B>A} = \frac{B_{ch\_A}}{B_{ch\_B}}$$

Pairwise compensation can now be solved with the following formula:

$$B_{comp} = \frac{D_{ch\_B} - (S_{A>B} \times D_{ch\_A})}{1 - (S_{A>B} \times S_{B>A})}$$

where $B_{comp}$ is the compensated intensity for channel B,

$D_{ch\_B}$ is the intensity detected in channel B, and

$D_{ch\_A}$ is the intensity detected in channel A.

Compensation involving more than two colors were performed by multiplying the intensities of all overlapping channels with an inverse matrix of the spillover coefficients of all overlapping channels[69].

Intensity attenuation along z-axis of each channel was then examined by plotting the mean intensity of each slice (thresholded above background) along the z-axis. In most cases, intensity attenuation was minimal across the entire 3D volume and no further correction was required. In some images where a significant intensity drop was observed, a reference z-slice was used and the intensity of each z-slice was multiplied to match the mean intensity of the reference slice. Further trimming of the z-stack was performed in rare cases where high non-specific staining was observed near the surface of the tissues and when antibody staining did not fully penetrate the entire 3D volume.

*Segmentation*

Stardist3D was used for segmentation of lymphoid-shaped cells. A modified version was developed to enable accelerated processing of large volumetric datasets on the NIH HPC Biowulf cluster.

First, a training dataset was annotated semi-automatically. Briefly, a small volume of an image containing a representative distribution of ~100-200 cells was cropped and preliminary segmentation was performed using seeded watershed 3D segmentation. The cell labels were then manually corrected in *napari* and used as annotated labels for training a custom Stardist3D model.

Nuclear Ki-67 staining was typically used as the segmentation channel, but for certain images, GFP+ fluorescent cells were segmented with the GFP channel, and CD45.1+ or CD45.2+ donor cells were segmented using a pseudo-nuclei channel created from the CD45.1/2 membrane stains. To generate the pseudo-nuclei channel, membrane stains were first labeled using the Pixel Classification workflow in *ilastik*[70] and the probability layer exported for further processing. A Python-based pipeline was used to enhance the contrast of the membrane layer. Briefly, a Gaussian smoothing filter of bandwidth σ=3 pixels was first applied to each z-slice, followed by morphological erosion (disk footprint with radius=3), white tophat (disk footprint, radius=3) and adaptive histogram equalization (kernel=(9,9)) from the *scikit-image* libraries. The intensity of this *contrast-enhanced membrane* layer was then inverted to generate the *pseudo-nuclei* layer. To remove the inverted background intensity, the *contrast-enhanced membrane* layer was further processed with morphological filtering to generate a *closed mask* layer of the CD45.1/2 cells. Inverted background intensity was then excluded by multiplying the *pseudo-nuclei* layer with the *closed mask* layer. These two layers (*contrast-enhanced membrane* layer and *pseudo-nuclei* layer), together with a manually annotated cell label layer, were used as a multi-channel training dataset for Stardist3D segmentation.

A new custom model was trained for each new set of imaging data. The prediction steps were then performed on NIH HPC Biowulf cluster with the full-sized segmentation channel. Briefly, training and prediction of the 3D U-Net model was performed on GPU nodes, whereas the CPU-intensive non-maximal suppression and labeling steps were processed on multi-purpose CPU nodes. The output segmentation image containing individual cell labels (nuclear masks) was then used for downstream processing.

*Segmentation mask generation and data extraction*

Segmentation labels, if trained on nuclear stains, will not contain the membrane and cytoplasmic regions of the cells. To enable staining intensity information to be extracted, a membrane/cytoplasmic cell mask was first generated through morphological filtering using the *scikit-image* library, with dilation (radius=6) and erosion (radius=3) of the initial nuclear mask, followed by subtraction of the eroded mask from the dilated mask to create a new membrane/cytoplasmic mask. Gaussian filtering was performed to apply a weighted gradient to the masks.

To obtain the voxel intensities of single cells, cell boundary coordinates of each segmented cell were used to extract a 3D patch of the image channels (a rectangular cuboid containing fluorescent markers of the single cell). This 3D patch was multiplied with the masks (membrane and/or nuclear) to retain only the voxels within the cell mask. The mean intensity of each fluorescent channel was obtained by dividing the summed voxel intensities by the sum of mask values. The final output is a data array containing the cell coordinates (x, y, z), mean marker intensities of each channel as output values. User-defined custom mathematical functions can also be added as a module to generate additional parameters if desired. Due to the large number of segmented cells, a computational pipeline optimized for distributed processing on HPC clusters was developed and deployed for routine operations.

*Histocytometric gating, quantification of protein expression and subset classification*

Raw data extracted based on cell masks and protein marker channels, together with their positional coordinates, were further analyzed using custom Python-based tools. Briefly, mean intensity of protein markers of single cells were projected on two-dimensional histocytometric plots visualized using *matplotlib* library, and gating was performed to select subsets of cells for further analyses. Activated donor OT-I cells were selected based on co-expression of Ki-67 and GFP/CD45.1. For polyclonal activated CD8+ T cells, the subset was gated based on Ki-67 and CD8b expression. To enhance the membrane staining signal from background noise, the ratio of mean intensity in the membrane mask (protein signal) versus the nuclear mask (background noise) was used to obtain a signal-to-noise ratio value. Alternatively, the mean nuclear intensity (background noise) of a membrane marker was subtracted from the mean membrane intensity (protein signal) to perform background subtraction. Negative values after the subtraction (indicating signal-to-noise ratio of <1) were set to zero.

Further subsets were generated based on TCF-1 and PD-1 expression to define PD-1$^{+}$ and PD-1$^{-}$ TCF-1$^{+}$ $T_{SL}$, as well as TCF-1$^{-}$ $T_{EFF}$. The spatial distribution of each subset was then visualized using Imaris 10.0 and XT extension (Bitplane) through creating new Spots layers of each subset. Each Spots layer corresponding to a subset was then set to a different color for visualization purpose.


69 Gerner, M. Y., Kastenmuller, W., Ifrim, I., Kabat, J. & Germain, R. N. Histo-cytometry: a method for highly multiplex quantitative tissue imaging analysis applied to dendritic cell subset microanatomy in lymph nodes. *Immunity* **37**, 364-376 (2012). https://doi.org/10.1016/j.immuni.2012.07.011

70 Berg, S. *et al.* ilastik: interactive machine learning for (bio)image analysis. *Nat Methods* **16**, 1226-1232 (2019). https://doi.org/10.1038/s41592-019-0582-9
